# Androgen receptor inhibition induces metabolic reprogramming and increased reliance on oxidative mitochondrial metabolism in prostate cancer

**DOI:** 10.1101/2022.05.31.494200

**Authors:** Preston D. Crowell, Jenna M. Giafaglione, Anthony E. Jones, Nicholas M. Nunley, Takao Hashimoto, Amelie M.L. Delcourt, Anton Petcherski, Matthew J. Bernard, Rong Rong Huang, Jin-Yih Low, Nedas Matulionis, Xiangnan Guan, Nora M. Navone, Joshi J. Alumkal, Michael C. Haffner, Huihui Ye, Amina Zoubeidi, Heather R. Christofk, Orian S. Shirihai, Ajit S. Divakaruni, Andrew S. Goldstein

## Abstract

Prostate cancer cells that survive clinical androgen receptor (AR) blockade mediate disease progression and lethality. Reprogrammed metabolic signaling is one mechanism by which tumor cells can survive treatment. However, how AR inhibition reprograms metabolism, and whether altered metabolism can be exploited to eradicate cells that survive AR blockade, remains unclear. Here, we comprehensively characterized the effect of AR blockade on prostate cancer metabolism using transcriptomics, metabolomics, and bioenergetics approaches. AR inhibition maintains oxidative mitochondrial metabolism and reduces glycolytic signaling, through hexokinase II downregulation and decreased MYC activity. Robust elongation of mitochondria via reduced DRP1 activity supports cell fitness after AR blockade. In addition, AR inhibition enhances sensitivity to complex I inhibitors in several models, suggesting that AR blockade increases reliance on oxidative mitochondrial metabolism. Our study provides an enhanced understanding of how AR inhibition alters metabolic signaling and highlights the potential of therapies that target metabolic vulnerabilities in AR-inhibited cells.

## Introduction

Prostate cancer is the leading cause of cancer-related death in non-smoking males in the United States^1^. Prostate cancer progression from localized to advanced metastatic disease is driven by aberrant androgen receptor activity. Therefore, patients with metastatic prostate cancer are treated with androgen deprivation therapies (ADTs), which dampen AR activity by depleting the levels of circulating androgens, alone or in combination with chemotherapy^2^. Prostate cancer that responds to ADT is termed castration-sensitive prostate cancer (CSPC). Prostate cancer that recurs after ADT is termed castration-resistant prostate cancer (CRPC)^3^. As AR activation remains critical for the survival and growth of the majority of CRPC cells, CRPC is treated with androgen-receptor pathway inhibitors (ARPIs), including enzalutamide which directly interacts with AR to impair its function^4^. Although ARPIs are initially effective, prolonged ARPI treatment invariably leads to treatment resistance and disease progression, ultimately causing lethality^5^. New approaches are needed to understand how prostate cancer cells survive ADT and/or ARPI treatment in order to target them and prevent or delay disease progression.

Prostate cancer initiation and progression are associated with metabolic reprogramming and several studies suggest that metabolic pathways can be targeted in prostate cancer to impair tumor growth^6-13^. For example, targeting lipogenesis via FASN inhibition or targeting glutamine metabolism via glutaminase inhibition antagonizes CRPC^7,8^. Additionally, CAMKK2 inhibition impairs CSPC and CRPC growth by disrupting autophagy^9,10^. Furthermore, serine biosynthesis activity and lactate export have been targeted to reduce growth in models of neuroendocrine prostate cancer^11,13^. While stimulation of AR signaling has been shown to promote anabolic metabolism, the effect of AR blockade on the metabolic signaling of prostate cancer cells has not been comprehensively defined. Therefore, it is critical to determine how metabolism is reprogrammed in the cells that survive AR inhibition in order to exploit therapy-induced metabolic vulnerabilities to delay or prevent prostate cancer progression.

In this study, we hypothesized that the metabolic requirements and vulnerabilities of AR-driven prostate cancer cells may shift as a result of AR inhibition. We utilized a variety of models and approaches to define how AR blockade alters the metabolic phenotype of prostate cancer cells. AR inhibition reprograms the metabolome in a consistent manner *in vitro* and *in vivo*. Cells that survive AR blockade are able to maintain oxidative mitochondrial metabolism while exhibiting reduced glycolysis driven by HK2 downregulation and decreased MYC activity. Mitochondrial elongation, via reduced DRP1-driven mitochondrial fission, enables AR inhibited cells to better survive AR blockade. We explored whether AR inhibition results in increased reliance on oxidative mitochondrial metabolism and observed enhanced sensitivity to complex I inhibitors after AR blockade. Taken together, our data suggest that AR blockade reprograms cellular metabolism and increases dependence on oxidative mitochondrial metabolism.

## Results

### Transcriptomic and metabolomic profiling reveal AR inhibition-induced metabolic reprogramming

To gain insight into how prostate cancer cells survive AR inhibition, we evaluated which pathways are altered after clinical AR blockade using the *Rajan et al* dataset^14^ which contains transcriptomics data from 7 patient tumors collected prior to and after androgen deprivation therapy (ADT). 10 of the top 30 significantly altered pathways identified by KEGG PATHWAY analysis were metabolism-related (Fig. 1a). To model transcriptional responses to extended AR inhibition, we treated the 16D CRPC cell line^15^ with 10μM enzalutamide for more than two months, termed LTenza for Long-Term enzalutamide-treatment. Gene Set Enrichment Analysis (GSEA) identified negative enrichment of Hallmark_androgen_response genes in LTenza 16D cells (Supplementary Figure 1a), validating AR inhibition. Transcriptomics analysis revealed that enzalutamide-naïve (vehicle-treated) 16D cells cluster with pre-ADT clinical samples, whereas LTenza 16D cells cluster with post-ADT samples from the *Rajan et al* dataset^14^ (Fig. 1b). 16D cells cultured with enzalutamide for up to 48 hours, termed STenza for Short-Term enzalutamide-treatment, clustered in between naïve 16D and LTenza 16D cells (Supplementary Figure 1b). Both STenza and LTenza 16D cells contained increased expression of genes upregulated post-ADT in the *Rajan et al* dataset, with LTenza cells containing the highest expression of such genes (Fig. 1c). Differential expression analysis identified 2074 enzalutamide-upregulated and 1498 enzalutamide-downregulated genes in LTenza 16D cells (Fig. 1d). KEGG PATHWAY analysis on the differentially expressed genes identified 12 metabolism-related pathways among the top 30 significantly altered pathways (Fig. 1e). Taken together, these data provide strong evidence that (1) AR inhibition modulates metabolic gene expression, and (2) enzalutamide treatment of 16D cells models transcriptional responses to clinical AR blockade.

**Figure 1.**
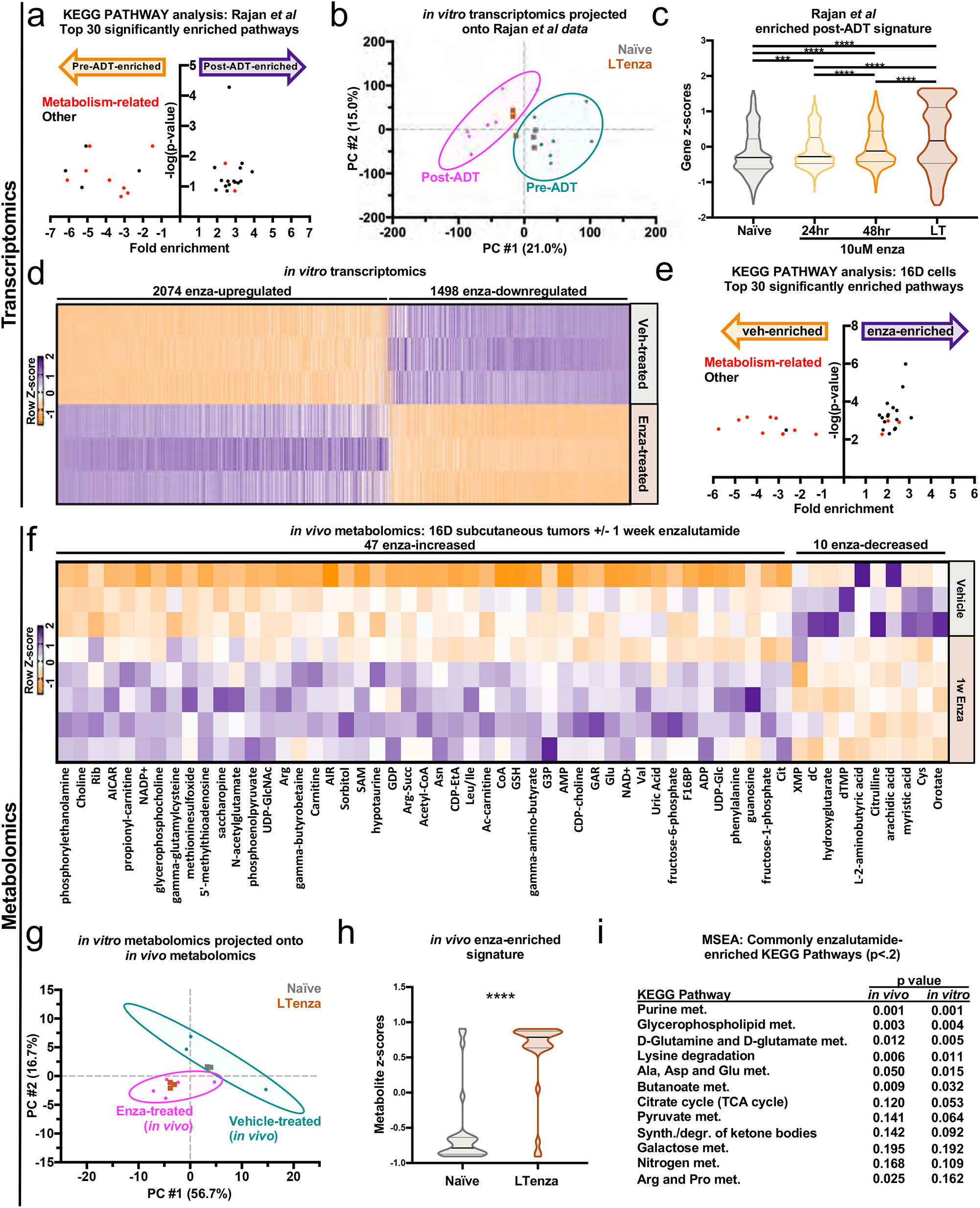
Transcriptomic and metabolic profiling identify AR-inhibition-induced metabolic reprogramming. (a) Top 30 significantly enriched pathways identified by KEGG PATHWAY analysis on differentially expressed (fold change ≥ 2, FDR < 0.2) *Rajan et al* pre-androgen deprivation therapy (Pre-ADT) and post-androgen deprivation therapy (Post-ADT) genes. Metabolism-related pathways highlighted in red. (b) Naïve and LTenza 16D transcriptomics data projected onto principle component analysis (PCA) plot of pre-ADT and post-ADT samples from *Rajan et al* data. 95% confidence eclipses for pre- and post-ADT data are shown in cyan and pink respectively. (c) Violin plot indicating gene z-scores of 1023 *Rajan et al* genes enriched post-ADT (fold change ≥ 2, FDR < 0.2) in naïve, 24hr enzalutamide-treated (enza), 48hr enza, and LTenza 16D cells. Data represent mean +/- SEM. (d) Heatmap of differentially expressed genes (fold change ≥ 2, FDR < 0.05) in LTenza 16D cells (Enza-treated) compared to naïve (Veh-treated) 16D cells. (e) Top 30 significantly enriched pathways identified by KEGG PATHWAY analysis on differentially expressed genes (fold change ≥ 2, FDR < 0.05) in naïve and LTenza 16D cells. Metabolism-related pathways highlighted in red. (f) Heatmap of differentially abundant metabolites (fold change ≥ 1.25, FDR < 0.2) in 1-week enzalutamide-treated 16D tumors compared to vehicle-treated 16D tumors. (g) *In vitro* naïve and LTenza 16D metabolomics data projected onto PCA plot of vehicle-treated and enza-treated samples from *in vivo 16D* metabolomics. 95% confidence eclipses for vehicle- and enza-treated *in vivo* data are shown in cyan and pink respectively. (h) Violin plot indicating metabolite z-scores of 47 *in vivo* enza-enriched metabolites (fold change ≥ 1.25, FDR < 0.2) in naïve and LTenza 16D cells. Data represent mean +/- SEM. (i) Metabolite Set Enrichment Analysis (MSEA) on *in vivo* enzalutamide-enriched metabolites (fold change ≥ 1.25, FDR < 0.2) and *in vitro* enzalutamide-enriched metabolites (fold change ≥ 1.25, FDR < 0.05) identifies commonly-enriched KEGG pathways (p<.2). P-values were calculated from an unpaired t-test with Welch’s correction (c and h) and a Fisher’s Exact Test (i). ***p < 0.001, ****p < 0.0001.

Having identified transcriptional evidence of AR blockade-induced metabolic reprogramming, we asked whether enzalutamide treatment of 16D cells alters the metabolome. NOD SCID IL2Rγ^null^ (NSG) mice bearing subcutaneous 16D tumors were treated with vehicle or enzalutamide for 10 days prior to tumor harvest, metabolite extraction, and metabolic profiling by high performance liquid chromatography mass spectrometry. Enzalutamide-treated tumors exhibited reduced protein expression of PSA, an AR target, and increased expression of NSE, which is repressed by AR, confirming AR inhibition *in vivo* (Supplementary Figure 1c). Metabolomic profiling of vehicle- and enzalutamide-treated tumors identified 47 enzalutamide-increased and 10 enzalutamide-decreased metabolites (Fig. 1f). We asked whether *in vitro* enzalutamide treatment similarly alters the metabolome by performing metabolic profiling on naïve and LTenza 16D cells. These analyses revealed that metabolic profiles group based on treatment, as naïve 16D cells cluster with vehicle-treated 16D tumors, whereas LTenza 16D cells cluster with enzalutamide-treated 16D tumors (Fig. 1g). In addition, we observed a higher abundance of *in vivo* enzalutamide-enriched metabolites in LTenza 16D cells compared to naïve 16D cells (Fig. 1h). *In vitro* metabolomics identified 32 enzalutamide-increased and 8 enzalutamide-decreased metabolites (Supplementary Figure 1d). To identify metabolic pathways commonly altered *in vivo* and *in vitro*, we performed Metabolite Set Enrichment Analysis (MSEA) on the enzalutamide-increased metabolites from each dataset. Among the commonly enriched KEGG pathways were terms related to lipid metabolism and the tricarboxylic acid cycle (TCA cycle) (Fig. 1i).

### AR blockade maintains oxidative phosphorylation and reduces glycolysis

Having identified enzalutamide-induced changes to the metabolome, we explored whether AR inhibition of 16D cells alters bioenergetics by measuring oxygen consumption rate (OCR) and extracellular acidification rate (ECAR) in naïve and LTenza 16D cells^16,17^ (Fig. 2a,b). Although enzalutamide treatment did not significantly alter ATP-linked respiration (Fig. 2a,c), FCCP-stimulated respiration was increased in enzalutamide-treated cells (Fig. 2a,d), demonstrating an enhanced maximal capacity for oxidative mitochondrial metabolism. We then transformed rates of OCR and ECAR into rates of mitochondrial and glycolytic ATP production to quantify the redistribution between oxidative phosphorylation and glycolysis upon enzalutamide treatment^18^. The mitochondrial ATP production rate was not altered in LTenza 16D cells (Fig. 2a,e), whereas the glycolytic ATP production rate was dramatically reduced (Fig. 2b,f). As such, the total ATP production rate in LTenza 16D cells (Fig. 2g) was substantially reduced, and oxidative phosphorylation comprised a greater percentage of the overall ATP supply (Fig. 2h). Consistent with dampened glycolysis, lower steady-state lactate was observed in LTenza 16D cells (Fig. 2i).

**Figure 2.**
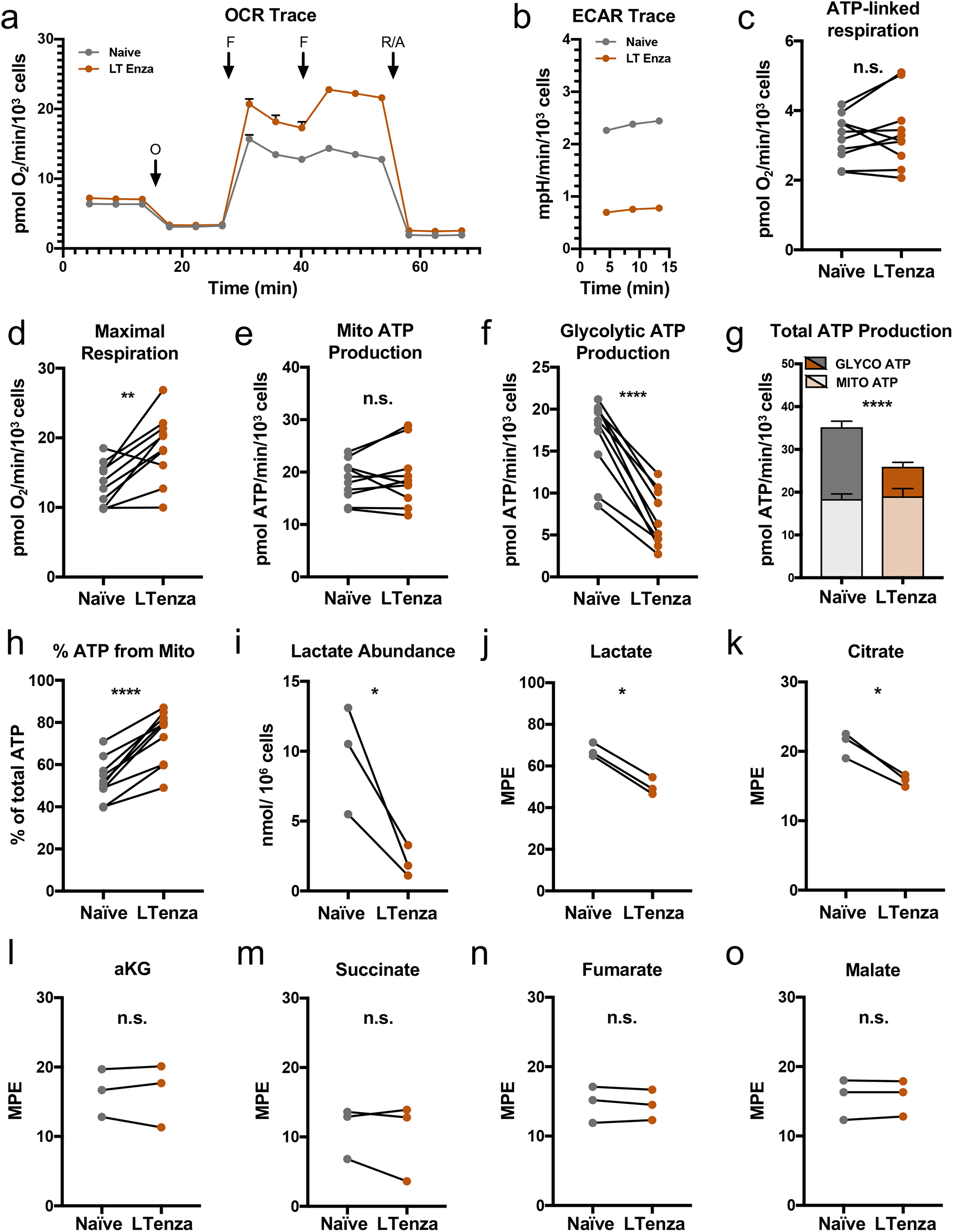
AR blockade maintains oxidative mitochondrial metabolism and reduces glycolysis. (a and b) Representative kinetic trace plots of the Oxygen Consumption Rate (OCR) (a) and Extracellular Acidification Rate (ECAR) (b) of naïve and LTenza 16D cells. Treatment with Oligomycin (O), FCCP (F), Rotenone and Antimycin A (R/A) are indicated with arrows. Data represent mean +/- SEM. (c and d) ATP-linked respiration (c) and maximal respiration (d) of naïve and LTenza 16D cells from 10 biological replicate experiments. (e and f) Mitochondrial (Mito) ATP production (e) and glycolytic ATP production (f) of naïve and LTenza 16D cells from 10 biological replicate experiments. (g) Total ATP production as the sum of mitochondrial ATP production (Mito ATP) and glycolytic ATP production (Glyco ATP) of naïve and LTenza 16D cells from 10 biological replicate experiments. Statistics refer to comparison of total ATP levels. Data represent mean + SEM. (h) Percentage of total ATP production from mitochondrial ATP production (% ATP from Mito) of naïve and LTenza 16D cells from 10 biological replicate experiments. (i) Lactate abundance in naïve and LTenza 16D cells from 3 biological replicate experiments. (j - o) Moles percent enrichment (MPE) of U-13C_6_-labeled glucose in lactate (j), citrate (k), alpha-ketoglutarate (aKG) (l), succinate (m), fumarate (n), and malate (o) in naïve and LTenza 16D cells from 3 biological replicate experiments. P-values were calculated from a ratio paired t-test. *p < 0.05, **p < 0.01, ****p < 0.0001, n.s. = not significant, p ≥ 0.05.

As a complementary approach to respirometry, we performed isotope tracing with U-13C_6_-labeled glucose to confirm a relative shift away from glycolysis and towards oxidative phosphorylation upon enzalutamide treatment. There was less glucose enrichment into lactate in LTenza cells, consistent with reduced glycolysis (Fig. 2j). Unlike the substantial decrease in labeling from glucose into lactate, enrichment into TCA cycle intermediates was largely maintained between naive and long-term enzalutamide-treated cells (Fig. 2k – o). The lone exception was a slight increase in relative flux from glucose into citrate in naïve cells (Fig. 2k), which could be indicative of decreased *de novo* lipogenesis upon AR inhibition^19^. Our data support a model whereby AR inhibition leads to reduced glycolysis but maintenance of oxidative mitochondrial metabolism. Interestingly, similar features have been reported in triple negative breast cancer cells that survive chemotherapy^20^, suggesting that LTenza cells may adopt a metabolic phenotype associated with treatment-resistance in other epithelial tumor types.

### Impaired MYC activity and downregulation of Hexokinase 2 contribute to AR inhibition-induced metabolic reprogramming

We wondered what mechanisms induced by AR blockade may contribute to the reduction in glycolysis. Transcriptomic analysis identified a trend toward downregulation of glycolytic genes in LTenza 16D cells (Fig. 3a). Among the most downregulated genes were *Hexokinase 2* (*HK2*) and *Lactate Dehydrogenase A* (*LDHA*) (Fig. 3a). Western blot analysis confirmed reduced protein expression of HK2 and LDHA in LTenza 16D cells (Fig. 3b). In addition, week-long enzalutamide-treated subcutaneous 16D tumors exhibited robust downregulation of HK2 and LDHA suggesting consistent enzalutamide-induced downregulation of HK2 and LDHA *in vivo* and *in vitro* (Fig. 3c). We performed immunohistochemistry on tissue sections from 16D tumors and observed relatively uniform HK2 downregulation in enzalutamide-treated tissues (Supplementary Figure 2a). To better understand the *in vivo* regulation of glycolytic enzymes following AR inhibition, we utilized an AR-positive patient-derived xenograft (PDX) model originating from a patient with localized CRPC, termed 180-30^21^. We confirmed reduced PSA and increased NSE expression in 1-week enzalutamide-treated 180-30 tumors (Fig. 3d). Consistent with our findings in the 16D model, enzalutamide-treated 180-30 tumors contained reduced expression of HK2 and LDHA (Fig. 3d).

**Figure 3.**
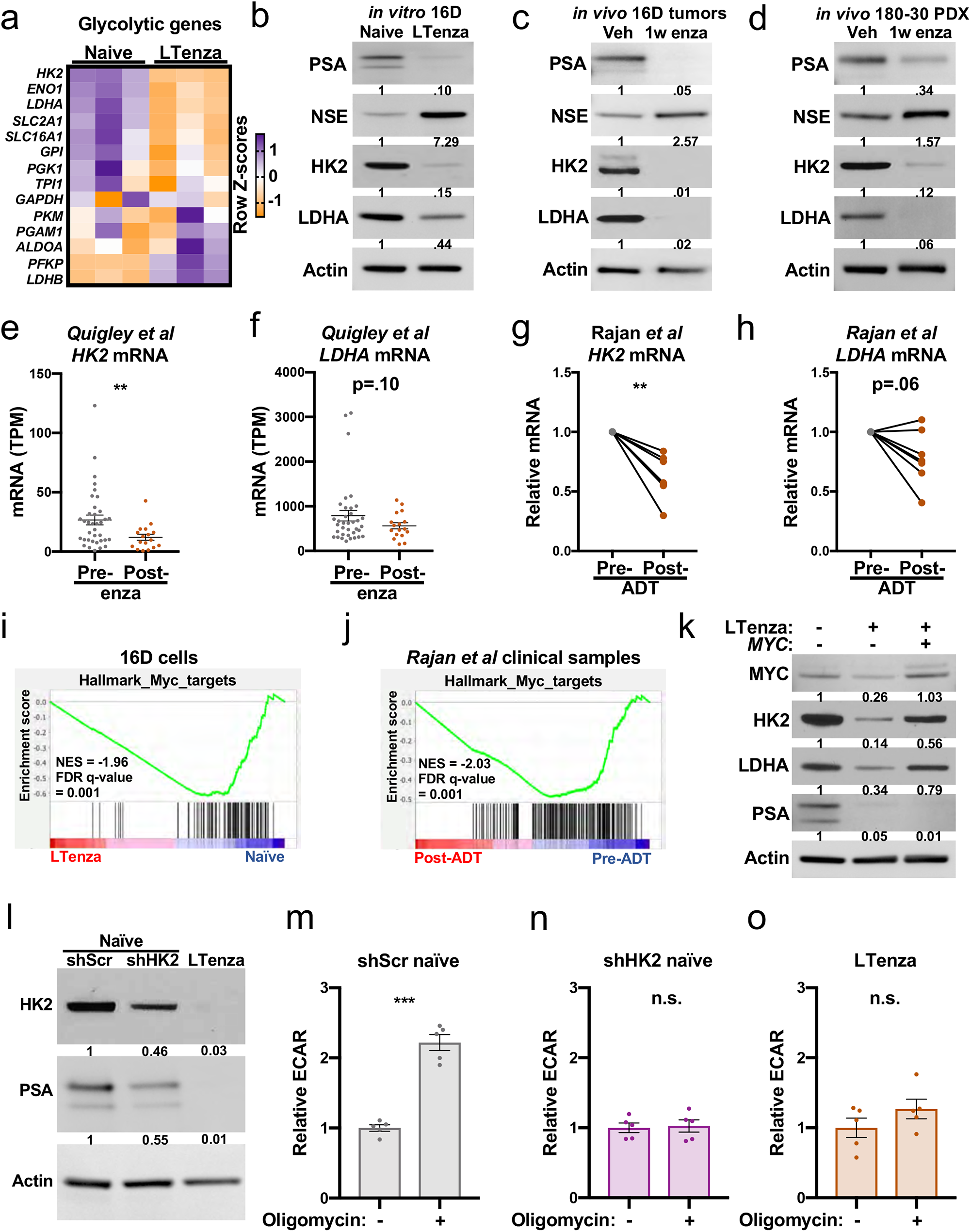
HK2 downregulation after AR inhibition contributes to reduced glycolytic capacity. (a) Heatmap showing the mRNA expression of glycolytic genes from RNA sequencing of 3 technical replicates of naïve and LTenza 16D cells. (b-d) Western blots indicating the expression of PSA, NSE, HK2, LDHA, and Actin (control) in lysates from naïve and LTenza 16D cells cultured *in vitro* (b), vehicle (veh) and 1-week enzalutamide-treated (1w enza) 16D tumors *in vivo* (c), and veh and 1w enza 180-30 patient derived xenografts *in vivo* (180-30 PDX) (d). (e and f) *HK2* (e) and *LDHA* (f) mRNA expression in unmatched enzalutamide-naïve (Pre-enza) and enzalutamide-treated (Post-enza) metastatic CRPC biopsies from *Quigley et al* dataset. Data represent the mean +/- SEM. (g and h) *HK2* (g) and *LDHA* (h) mRNA expression in matched pre- and post-androgen deprivation therapy (ADT) biopsies from the *Rajan et al* dataset. (i and j) GSEA of Hallmark_Myc_targets in naïve and LTenza 16D cells (i), and *Rajan et al* pre-ADT and post-ADT samples (j) showing normalized enrichment scores (NES) and false discovery rates (FDR). (k) Western blot indicating expression of MYC, HK2, LDHA, PSA, and Actin (control) in naïve (-/-), RFP-transduced LTenza (+/-), and MYC-transduced (+/+) LTenza 16D lysates. (l) Western blot detecting HK2, PSA and Actin (control) in shScr-transduced naïve, shHK2-transduced naïve, and LTenza 16D lysates. (m - o) Relative Extracellular Acidification Rate (ECAR) of shScr-transduced naïve (m), shHK2-transduced naïve (n), and LTenza 16D cells (o) treated +/- Oligomycin. Data represent the mean +/- SEM of 5 technical replicates from a representative experiment (n=2). P-values were calculated from an unpaired t-test with Welch’s correction (e, f, m, n, and o) and a ratio paired t-test (g and h). **p < 0.01, ***p < 0.001, n.s. = not significant, p ≥ 0.05.

We next asked whether there is evidence of *HK2* and *LDHA* downregulation in clinical datasets after AR inhibition. Transcriptomics analysis of the *Quigley et al* dataset^22^, which contains enzalutamide-naïve and enzalutamide-treated metastatic CRPC biopsies, revealed significant *HK2* mRNA downregulation and a trend toward reduced *LDHA* levels in the enzalutamide-treated samples (Fig. 3e,f). To investigate whether AR inhibition-induced HK2 and LDHA downregulation is unique to CRPC or if it is broadly associated with AR blockade, we evaluated *HK2* and *LDHA* levels in castration-sensitive tumors before and after ADT using the *Rajan et al* dataset^14^. *HK2* mRNA expression was reduced in all 7 patients post-ADT (Fig. 3g) while *LDHA* expression was reduced in 5 of 7 patients (Fig. 3h). These data suggest that AR inhibition lowers *HK2* and *LDHA* levels across various disease states.

Previous work suggests that AR can regulate select glycolytic genes^9^. We analyzed our previous AR ChIP-seq datasets in 16D cells^23^ and did not observe evidence of binding to the *HK2* or *LDHA* loci, suggesting that these glycolytic enzymes are not direct targets of AR in 16D cells. To explore an indirect mechanism of *HK2* and *LDHA* downregulation after AR inhibition, we investigated whether AR blockade alters transcriptional signatures of MYC, a key regulator of glycolysis^24^. GSEA revealed negative enrichment of Hallmark_Myc_targets in LTenza 16D cells (Fig. 3i). Consistent with these findings, we observed negative enrichment of Hallmark_Myc_targets in *Rajan et al* patient samples post-ADT^14^ (Fig. 3j). In addition, negative enrichment of Hallmark_Myc_targets was observed after castration in the AR positive LTL331 PDX model^25^ (Supplementary Figure 2b).

To determine whether reduced MYC activity mediates reduced HK2 and LDHA expression in AR inhibited cells, we attempted to rescue MYC activity via ectopic MYC expression in LTenza 16D cells. GSEA revealed positive enrichment of Hallmark_Myc_targets in MYC-transduced LTenza cells compared to RFP-transduced LTenza 16D cells (Supplementary Figure 2c). Furthermore, there was no significant negative enrichment of Hallmark_Myc_targets between MYC-transduced LTenza cells and naïve 16D cells indicating successful rescue of MYC transcriptional activity (Supplementary Figure 2d). Western blot analysis revealed increased expression of HK2 and LDHA in MYC-transduced LTenza cells compared to RFP-transduced LTenza 16D cells. However, HK2 expression remained roughly 50 percent lower in MYC-transduced LTenza cells than in naïve 16D cells (Fig 3k). Targeted bisulfite sequencing identified a statistically significant increase in the mean percentage of methylated CpGs within the transcriptional start site of *HK2* in LTenza 16D cells suggesting that epigenetic alterations may cooperate with reduced MYC activity to antagonize HK2 expression (Supplementary Figure 2e).

Since HK2 is upstream of LDHA and catalyzes the first committed step of glycolysis, we evaluated whether HK2 knockdown (shHK2) in naïve 16D cells is sufficient to reduce glycolytic activity, compared to a scrambled control shRNA (shScr). HK2 knockdown was confirmed by Western blot (Fig. 3l). To broadly measure cellular glycolytic capacity, we measured ECAR after treatment with the ATP synthase inhibitor oligomycin, which will stimulate an increase in glycolysis due to the loss of oxidative phosphorylation. Whereas oligomycin stimulated a 2-fold increase in ECAR of shScr-transduced 16D cells (Fig. 3m), the rate was unchanged in shHK2-transduced 16D cells (Fig. 3n) or LTenza 16D cells (Fig. 3o). Accordingly, shHK2-transduced 16D cells and LTenza 16D cells exhibited reduced oligomycin-stimulated ECAR compared to shScr-transduced 16D cells (Supplementary Figure 2f). These data establish reduced HK2 as one mechanism contributing to a lower glycolytic capacity in response to AR inhibition.

### Enzalutamide induces mitochondrial elongation via reduced DRP1 activity

As mitochondrial dynamics can change in response to cellular and environmental stresses^26^, we explored the effect of AR inhibition on mitochondrial morphology. Mitochondria were visualized in naïve and LTenza 16D cells by staining for the mitochondria-associated protein TUFM. Immunofluorescence identified robustly elongated mitochondria in LTenza 16D cells (Fig. 4a). Quantification of mitochondrial elongation and branching was performed by calculating the mitochondrial aspect ratio, which is equal to the ratio of the major axis to the minor axis of an object, and form factor, a value that compensates for irregularity in the shape of an object, respectively^27^ (Fig. 4b). LTenza 16D cell mitochondria exhibited a higher aspect ratio (Fig. 4c) and lower form factor (Fig. 4d) compared to naïve cell mitochondria, consistent with mitochondrial elongation and increased branching. Eccentricity, the ratio of the distance between the foci of an ellipse and its major axis length, was used as a secondary approach to quantify mitochondrial elongation. Increased mitochondrial eccentricity was calculated in LTenza 16D cells, confirming mitochondrial elongation (Supplementary Figure 3a). Enzalutamide treatment did not alter mitochondrial size, subtly increased mitochondrial count, and did not alter mitochondrial volume (Supplementary Figure 3b-d). As mitochondrial elongation and branching have been associated with enhanced mitochondrial function in certain contexts^28^, these features may enable cells to compensate for reduced glycolytic activity after AR inhibition.

**Figure 4.**
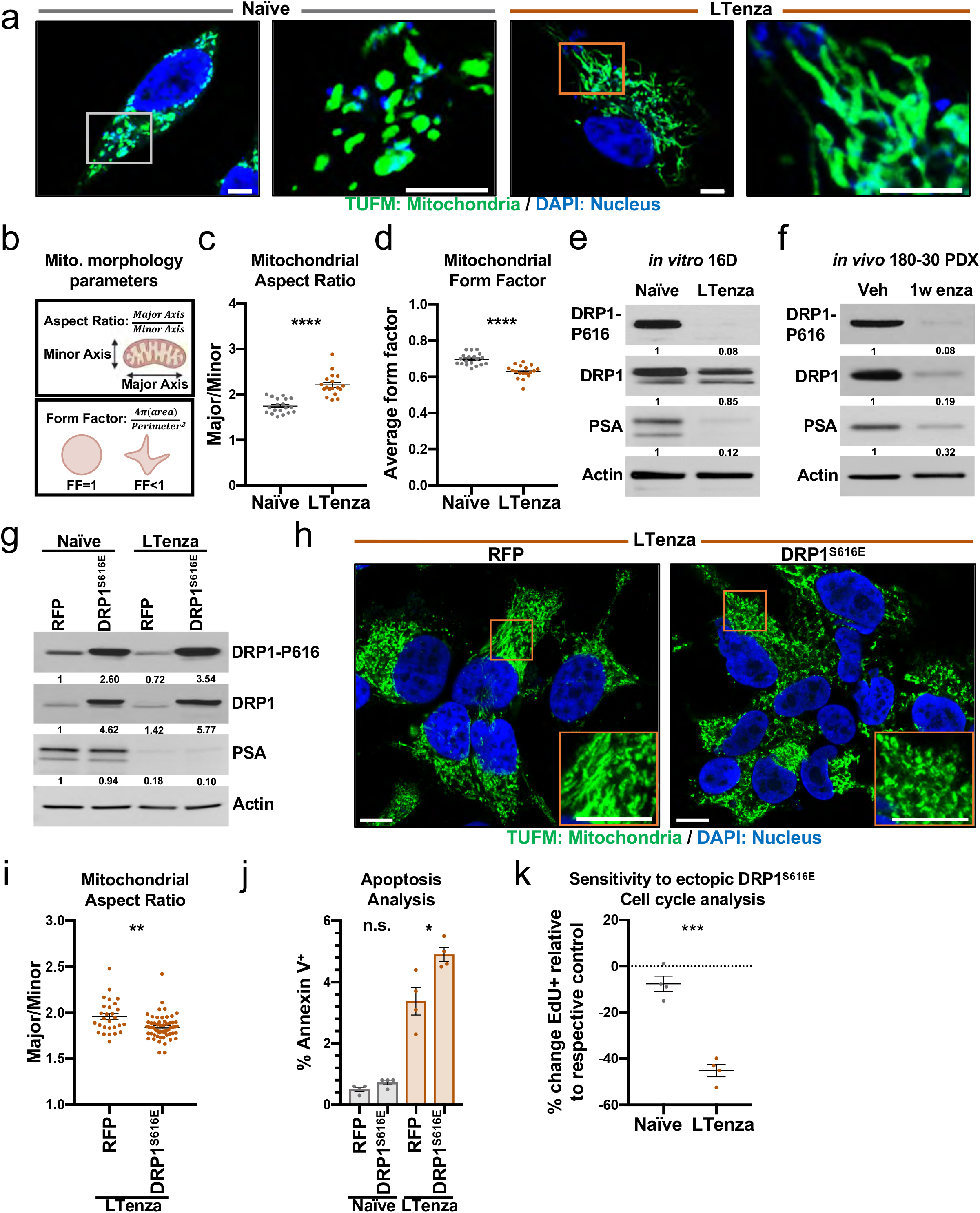
AR blockade elongates mitochondria via reduced DRP1 activity. (a) Representative immunofluorescent images of naïve and LTenza 16D cells stained for TUFM (green) and DAPI (blue). Scale bars, 5 μm. (b) Schematic illustrating calculation of aspect ratio and form factor. (c and d) Quantification of mitochondrial aspect ratio (c) and mitochondrial form factor (d) from TUFM stains from 20 images per treatment group. Data represent the mean +/- SEM. (e and f) Western blots detecting DRP1 phosphorylation at S616 (DRP1-P616), DRP1, PSA, and Actin (control) in naïve and LTenza 16D lysates (e), and vehicle (veh) and 1-week enzalutamide-treated (1w enza) 180-30 PDX tumor lysates (f). (g) Western blot indicating DRP1-P616, DRP1, PSA, and Actin (control) expression in lysates from RFP- or DRP1^S616E^-transduced naïve and LTenza 16D cells. (h) Representative immunofluorescent images of RFP- and DRP1^S616E^-transduced LTenza 16D cells stained for TUFM (green) and DAPI (blue). Scale bars, 10 μm. (i) Quantification of mitochondrial aspect ratio from TUFM stains from at least 28 cells per treatment group. Data represent the mean +/- SEM. (j) Apoptosis analysis to identify the percentage of Annexin V-positive cells (% Annexin V^+^) in each transduced line after 48 hours of culture. Data represent the mean +/- SEM of 4 technical replicates. (k) Cell cycle analysis to quantify the relative sensitivity of naïve and LTenza 16D cells to ectopic DRP1^S616E^ expression. Data represent the mean +/- SEM of 4 technical replicates from a representative experiment (n=2). P-values were calculated from an unpaired t-test with Welch’s correction. *p < 0.05, **p < 0.01, ***p < 0.001, ****p < 0.0001, n.s. = not significant, p ≥ 0.05.

Mitochondrial morphology is determined by the relative amounts of mitochondrial fission and fusion^29^. Several reports provide evidence that AR may regulate DRP1^30^, encoded by the *DNM1L* gene, which mediates mitochondrial fission. We therefore explored whether DRP1 levels are altered in LTenza 16D cells. DRP1 expression was only subtly reduced in LTenza 16D cells (Fig. 4e). As DRP1 phosphorylation at S616 is required for DRP1 activity^31^, we hypothesized that LTenza 16D cells may exhibit reduced DRP1-S616 phosphorylation. Indeed, DRP1-S616 phosphorylation was dramatically reduced in LTenza cells compared to naïve 16D cells (Fig. 4e). Enzalutamide-treated 16D tumors contained both reduced total DRP1 expression and reduced DRP1-S616 phosphorylation (Supplementary Figure 3e,f). Furthermore, we observed both reduced total DRP1 expression and reduced DRP1-S616 phosphorylation in 180-30 PDX tumors suggesting that the tumor microenvironment may influence the response of DRP1 expression to AR blockade (Figure 4f). Analysis of 16D AR ChIP-seq data^23^ revealed binding to the *DNM1L* locus, suggesting that AR directly regulates DRP1 in this model (Supplementary Figure 3g).

To evaluate the functional role of DRP1, we ectopically expressed a constitutively active DRP1 phosphomimetic^23^, DRP1^S616E^, in naïve and LTenza 16D cells (Fig. 4g). DRP1^S616E^-transduced LTenza 16D cells contained more fragmented mitochondria than RFP-transduced LTenza 16D cells (Fig. 4h,i). Apoptosis and cell cycle analyses revealed that whereas naïve 16D cells are relatively insensitive to DRP1^S616E^ expression, DRP1^S616E^-transduced LTenza cells exhibit increased apoptosis and reduced proliferation compared to RFP-transduced LTenza cells (Fig. 4j,k). These data suggest that elongation of mitochondria may enable LTenza 16D cells to better survive enzalutamide treatment. Our data are consistent with previous reports that mitochondrial elongation can promote tumor cell survival during energetic stress^26,31,32^.

### AR inhibition enhances sensitivity to complex I inhibitors

As enzalutamide-treated 16D cells generate a greater proportion of ATP from oxidative mitochondrial metabolism and exhibit reduced glycolytic activity and higher respiratory capacity, we hypothesized that these cells may be increasingly sensitive to inhibition of oxidative phosphorylation. To test our hypothesis, we treated naïve and LTenza 16D cells with the highly-specific complex I inhibitor IACS-010759^33^ (IACS). Respirometry and U-13C_6_-glucose tracer analysis were performed to validate the on-target effect of IACS. IACS reduced the ATP-linked respiration of naïve and LTenza 16D cells by roughly 95 percent (Fig. 5a). In addition, IACS treatment significantly reduced M+2 labeling of TCA cycle intermediates from U-13C_6_-glucose in both naïve and LTenza 16D cells (Supplementary Figure 4a,b). Increased M+3-labeled lactate was observed in both naïve and LTenza 16D cells after IACS treatment indicating that both cell types compensate for reduced complex I activity by increasing glycolysis (Supplementary Figure 4c). Respirometry revealed that while both naïve and LTenza 16D cells increase glycolytic ATP production in response to IACS, naïve cells contain a 2-fold higher IACS-induced glycolytic ATP production rate compared to LTenza 16D cells (Supplementary Figure 4d). Accordingly, IACS treatment reduced the total ATP production of naïve cells by just 12% compared to a 29% reduction of total ATP production in LTenza 16D cells (Supplementary Figure 4e).

**Figure 5.**
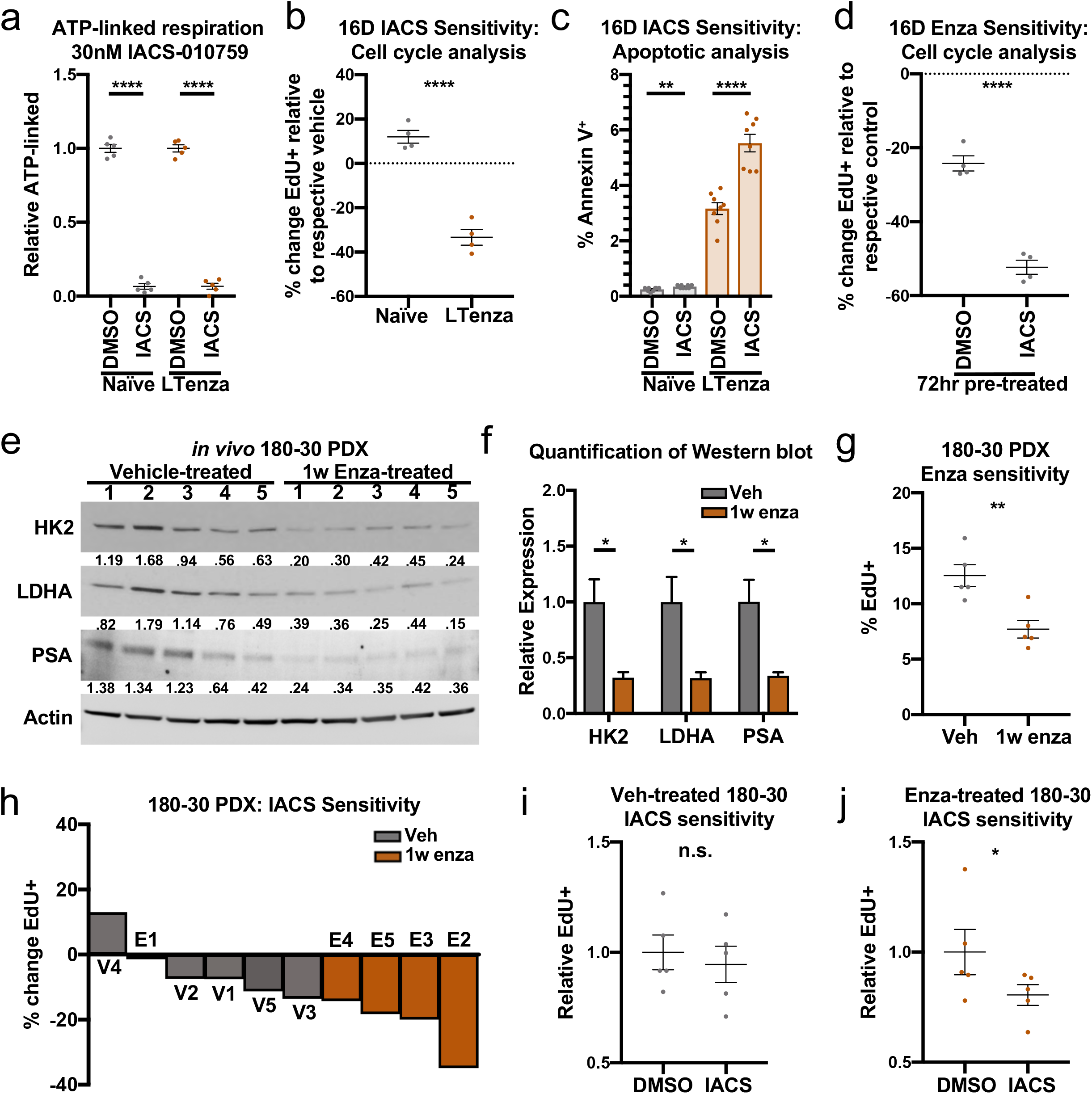
AR blockade enhances sensitivity to complex I inhibition. (a) ATP-linked respiration of naïve and LTenza 16D cells treated with vehicle (DMSO) or 30nM IACS-010759 (IACS) for 24 hrs. Data represent the mean +/- SEM of 5 technical replicates. (b) Cell cycle analysis to quantify the relative sensitivity of naïve and LTenza 16D cells to 30nM IACS. Data represent the mean +/- SEM of 4 technical replicates from a representative experiment (n=3). (c) Apoptosis analysis to identify the percentage of Annexin V-positive cells (% Annexin V^+^) in naïve and LTenza 16D cells treated with DMSO or 30nM IACS for 48 hours. Data represent the mean +/- SEM of 8 technical replicates. (d) Cell cycle analysis to quantify the relative sensitivity of DMSO- and 72hr 30nM IACS-treated naïve 16D cells to enzalutamide. Data represent the mean +/- SEM of 4 technical replicates. (e and f) Western blot detecting HK2, LDHA, PSA, and Actin (control) expression in lysates from 5 vehicle-treated and 5 1-week enzalutamide-treated 180-30 PDX tumors (e) and associated quantification (f). Data represent the mean +/- SEM. (g) Cell cycle analysis to quantify the proliferation (% EdU^+^) of vehicle-treated (veh) and 1w enza-treated 180-30 tumors after 3-day *ex vivo* culture in organoid conditions. Data represent the mean +/- SEM of 5 tumor samples per treatment group. (h) Waterfall plot indicating the *ex vivo* sensitivity of 180-30 PDX tumor tissue from veh- and 1w enza-treated tumors to 30nM IACS. Data represent the percent change in EdU positivity (% change EdU^+^) relative to the respective vehicle. (i and j) Cell cycle analysis of the sensitivity of vehicle-treated (i) or enza-treated (j) 180-30 PDX tumor tissue to *ex vivo* culture +/- 30nM IACS. Data represent the mean +/- SEM of 5 tumor samples per treatment group. P-values were calculated from an unpaired t-test with Welch’s correction (a-d, f and g) and a ratio paired t-test (i and j). *p < 0.05, **p < 0.01, ****p < 0.0001, n.s. = not significant, p ≥ 0.05.

We performed cell cycle analysis to determine the effect of IACS on the proliferation of naïve and LTenza 16D cells and identified robust differential sensitivity (Fig. 5b). Whereas IACS treatment did not alter the proliferation of naïve cells, IACS reduced the proliferation of LTenza 16D cells by roughly 35 percent in just 72 hours (Fig. 5b and Supplementary Figure 4f). Apoptosis analysis revealed that while IACS treatment increased apoptosis in both naïve and LTenza 16D cells, the percentage of apoptotic naïve cells remained far below one percent after 72 hours (Fig. 5c). In contrast, nearly 6% of IACS-treated LTenza cells were apoptotic after the same period (Fig. 5c). Since AR blockade increases sensitivity to complex I inhibition, we wondered if IACS treatment of naïve 16D cells might increase enzalutamide sensitivity. 72hr IACS pretreatment significantly enhanced enzalutamide sensitivity, effectively doubling the growth inhibition caused by enzalutamide (Fig. 5d). These data demonstrate the potential for combining IACS and enzalutamide to reduce prostate cancer cell proliferation regardless of which treatment is initiated first.

We explored whether the clinically viable drug metformin, which has complex I inhibitor activity *in vitro*^34^, alters the enzalutamide sensitivity of naïve 16D cells. Reduced ATP-linked respiration in metformin-treated 16D cells was confirmed by respirometry (Supplementary Figure 4g). Cell cycle analysis revealed that, unlike IACS, metformin alone was sufficient to impair the proliferation of naïve 16D cells (Supplementary Figure 4h). Reduced proliferation in metformin-treated 16D cells was likely caused by known off-target effects^34^ as IACS treatment reduced ATP-linked respiration by greater than 95% without altering EdU labeling. Consistent with IACS pretreatment increasing responsiveness to AR inhibition, 72hr metformin pretreatment significantly enhanced the sensitivity of naïve 16D cells to enzalutamide (Supplementary Figure 4i).

To better understand the interaction between AR inhibition and complex I inhibition across various disease states, we explored whether metformin similarly enhances the sensitivity of LNCaP cells to castration. After validating castration-induced suppression of AR target gene expression (Supplementary Figure 5a), and demonstrating that castrated LNCaP cells transcriptionally resemble castrated LTL331 PDX tumors^25^ and patient tumors post-ADT^14^ (Supplementary Figure 5b-e), we explored whether castrated LNCaP cells exhibit altered metabolic gene expression. GSEA revealed negative enrichment of Hallmark_Myc_targets (Supplementary Figure 5f), and we identified mRNA downregulation of select glycolytic genes including *HK2* and *LDHA* after castration (Supplementary Figure 5g). Western blot analysis confirmed lower HK2 and LDHA levels and identified reduced total DRP1 expression and DRP1-S616 phosphorylation (Supplementary Figure 5h). Consistent with our findings in the 16D model, metformin treatment reduced the growth of LNCaP cells and significantly increased castration sensitivity, from roughly 45 percent to greater than 85 percent (Supplementary Figure 5i,j). These data suggest that AR blockade-induced metabolic changes may be conserved across various disease states and that complex I inhibition could broadly enhance sensitivity to AR inhibition.

After showing that various complex I inhibitors can synergize with AR blockade *in vitro*, we explored whether enzalutamide treatment of mice bearing 180-30 PDX tumors enhances IACS sensitivity. After one week of treatment with vehicle or enzalutamide *in vivo*, we analyzed IACS sensitivity using *ex vivo* culture. Week-long enzalutamide-treated 180-30 PDX tumors contained reduced protein expression of PSA, HK2, and LDHA (Fig. 5e,f). Cell cycle analysis after *ex vivo* culture of tumor tissue in prostate organoid conditions confirmed reduced proliferation in enzalutamide-treated 180-30 PDX tumors (Fig. 5g). Analysis of IACS sensitivity revealed that enzalutamide-treated samples accounted for 4 of the 5 most IACS-sensitive samples (Fig. 5h). Furthermore, whereas IACS did not alter the growth of vehicle-treated tumor cells in a statistically significant manner (Fig. 5i), IACS significantly reduced proliferation of enzalutamide-treated cells (Fig. 5j).

## Discussion

Therapy-induced metabolic reprogramming has been reported in various cancers where standard of care therapy can synergize with targeting of reprogrammed metabolism to impair treatment-resistance^20,35,36^. In this study, we comprehensively characterized the effect of AR blockade on prostate cancer metabolism. Transcriptomic and metabolomic analyses revealed AR-inhibition-induced changes to metabolic gene expression and metabolite abundance respectively. We identified sustained oxidative mitochondrial metabolism, including increased maximal respiration, and reduced basal and oligomycin-stimulated glycolysis, after AR inhibition. Mechanistically, decreased MYC activity and HK2 downregulation contributed to reduced glycolytic activity. Interestingly, we observed robust elongation of mitochondria driven by lower DRP1 activity in enzalutamide-treated cells and found that mitochondrial elongation supports cell survival and proliferation after AR blockade. We explored whether AR inhibition increases reliance on oxidative mitochondrial metabolism and observed increased sensitivity to complex I inhibitors after AR blockade. In addition, pre-treatment with complex I inhibitors increased sensitivity to AR inhibition, demonstrating the effectiveness of combined complex I inhibition and AR blockade.

Our study identifies mitochondrial elongation as a potential survival mechanism after AR blockade. Elongation has been shown to protect mitochondria from autophagosomal degradation during nutrient starvation^26,32^. A hyperfused mitochondrial phenotype has been observed in triple negative breast cancer cells that survive chemotherapy^37^ and has been shown to enable chemotherapy resistance in gynecological cancers^38^. The functional impact of mitochondrial morphology on metabolic output has proven to be highly context dependent. For example, mitochondrial fission drives increased oxidative mitochondrial metabolism and tumorigenic potential in pancreatic ductal adenocarcinoma cells^39,40^ and decreased oxidative mitochondrial metabolism in neuroblastoma cells^41^. Future work is needed to understand how mitochondrial elongation enables prostate cancer cells to better survive AR blockade.

It is critical to consider the window when complex I inhibition could cooperate with AR inhibition to improve patient outcomes. Our functional data suggest that complex I inhibition could be combined with ADT to treat castration-sensitive prostate cancer and/or ARPIs to treat CRPC. These findings are further supported by our observation from clinical datasets that *HK2* downregulation occurs after both ADT treatment of localized castration-sensitive disease, and ARPI treatment of metastatic CRPC. While most prostate tumors initially respond to AR inhibition, they eventually recur in a more aggressive form, driven by the acquisition of additional somatic mutations such as disruption of RB1 and TP53^42,43^. Interestingly, knockdown of *RB1* and *TP53* in LTenza 16D cells was not sufficient to alter enzalutamide-induced metabolic features including reduced MYC activity and enhanced sensitivity to complex I inhibitors despite increased neuroendocrine signatures^44-46^ (Supplementary Figure 6). In contrast, in the LTL331 PDX model^25^, which relapses as terminally differentiated neuroendocrine prostate cancer after castration, relapsed tumors contained robust enrichment of Hallmark_Myc_targets compared to castrated tumors despite maintenance of low AR activity (Supplementary Figure 7). These data suggest that rescue of AR activity is not necessary to restore MYC signaling and that the effectiveness of combined AR blockade and complex I inhibitor treatment in relapsed ARPI-resistant disease may depend on the tumor phenotype and/or genetic driver. Future work is necessary to determine whether MYC reactivation is broadly associated with relapse, and whether increased MYC activity may contribute to prostate cancer recurrence by restoring glycolytic activity.

Metformin has been explored as a prostate cancer treatment for functions distinct from complex I inhibition. Specifically, metformin has been shown to inhibit the proliferation of prostate cancer cell lines *in vitro* by reducing AR and cyclin D1 levels^47,48^. These effects may explain why naïve 16D cells exhibit sensitivity to metformin, despite lacking sensitivity to IACS, which reduces mitochondrial respiration by greater than 95 percent. In addition, metformin has been shown to synergize with bicalutamide in mouse models by preventing AR blockade-induced hyperinsulinemia, which enhances tumor growth^49^. Accordingly, several observational and clinical trials have been performed and others are ongoing to determine the efficacy of combined metformin treatment and AR blockade^50^. Such trials have thus far been inconclusive regarding metformin use and both recurrence-free survival and overall survival^51^. Importantly, the concentration of metformin required to inhibit complex I activity *in vitro* (1mM) is more than 10 times higher than the maximally-achievable therapeutic concentration (70μM) found in patients^34,52^. Therefore, improved clinically-viable inhibitors of complex I are needed to evaluate the efficacy of combined AR blockade and inhibition of mitochondrial oxidation in patients.

## Acknowledgments

P.D.C. and J.M.G. acknowledge the support of the UCLA Eli and Edythe Broad Center of Regenerative Medicine and Stem Cell Research Training Program. P.D.C. is also supported by the NIH grants TL1 DK132768 and U2C DK129496. A.E.J. is supported by the UCLA Tumor Cell Biology Training Grant (NIH T32CA009056). A.S.G. is supported by the National Cancer Institute of the National Institutes of Health under Award Number R01CA237191. The content is solely the responsibility of the authors and does not necessarily represent the official views of the National Institutes of Health. A.M.L.D. is supported by NCI/NIH supplement related to R01CA237191. A.S.G. is also supported by American Cancer Society award RSG-17-068-01-TBG, the UCLA Eli and Edythe Broad Center of Regenerative Medicine and Stem Cell Research Rose Hills Foundation Innovator Grant, the UCLA Jonsson Comprehensive Cancer Center and Eli and Edythe Broad Center of Regenerative Medicine and Stem Cell Research Ablon Scholars Program, the National Center for Advancing Translational Sciences UCLA CTSI Grant UL1TR001881, STOP CANCER, and the UCLA Institute of Urologic Oncology. A.S.G., A.S.D. and O.S.S. are supported by the UCLA Prostate Cancer Specialized Programs of Research Excellence (SPORE) NCI P50 CA092131. J.J.A is supported by the National Cancer Institute award R01CA251245, the Pacific Northwest Prostate Cancer Specialized Programs of Research Excellence (SPORE) NCI P50 CA097186, the NCI Drug Resistance and Sensitivity Network NCI P50 CA186786-07S1, and the Department of Defense Idea Award (W81XWH-20-1-0405). M.C.H. is supported by the U.S. Department of Defense Prostate Cancer Research Program (W81XWH-20-1-0111, W81XWH-21-1-0229) and Grant 2021184 from the Doris Duke Charitable Foundation. We thank UCLA Technology Center for Genomics & Bioinformatics, UCLA Metabolomics Center, UCLA MCDB/BSCRC Microscopy Core. N.M.N. is supported by NCI U01 CA224044-03. The UCLA Integrated Technologies Core is supported by CURE/P30 DK041301. We thank Michaela Veliova, Thomas Graeber, Linsey Stiles, Matthew Rettig, Brigitte Gomperts, Paul Boutros and other collaborators who provided intellectual support and critical feedback during the course of this project.

## Author Contributions

P.D.C., J.M.G., A.E.J., N.M.N (Nunley), T.H., A.M.L.D., and M.J.B. conducted the experiments. P.D.C., A.E.J., A.S.D and A.S.G. designed the experiments, wrote and edited the manuscript. R.R.H. and H.Y. performed immunohistochemistry and provided pathology expertise and wrote the related methods section. A.P. and O.S.S. performed mitochondrial morphology quantification, provided metabolism expertise, and wrote the related methods section. J.L. and M.C.H. performed the DNA methylation analysis, and wrote the related methods section. N.M. and H.R.C. performed mass spectrometry on in vivo tumor metabolites, provided metabolism expertise and wrote the related methods section. X.G. and J.J.A. provided transformed data from the *Quigley et al* dataset and wrote the related methods section. A.Z. provided ChIP sequencing data, key cell lines, and wrote the related methods section. N.M.N (Navone) provided PDX models. A.S.G. procured funding and supervised the experiments.

## References

1. Siegel RL, Miller KD, Fuchs HE, Jemal A. Cancer statistics, 2022. CA Cancer J Clin. 2022;72(1):7–33.

2. Sayegh N, Swami U, Agarwal N. Recent Advances in the Management of Metastatic Prostate Cancer. JCO Oncol Pract. 2022;18(1):45–55.

3. Nakazawa M, Paller C, Kyprianou N. Mechanisms of Therapeutic Resistance in Prostate Cancer. Curr Oncol Rep. 2017;19(2):13. PMC5812366

4. Tran C, Ouk S, Clegg NJ, Chen Y, Watson PA, Arora V, Wongvipat J, Smith-Jones PM, Yoo D, Kwon A, Wasielewska T, Welsbie D, Chen CD, Higano CS, Beer TM, Hung DT, Scher HI, Jung ME, Sawyers CL. Development of a second-generation antiandrogen for treatment of advanced prostate cancer. Science. 2009;324(5928):787-790. PMC2981508

5. Schmidt KT, Huitema ADR, Chau CH, Figg WD. Resistance to second-generation androgen receptor antagonists in prostate cancer. Nat Rev Urol. 2021;18(4):209–226.

6. Priolo C, Pyne S, Rose J, Regan ER, Zadra G, Photopoulos C, Cacciatore S, Schultz D, Scaglia N, McDunn J, De Marzo AM, Loda M. AKT1 and MYC induce distinctive metabolic fingerprints in human prostate cancer. Cancer Res. 2014;74(24):7198–7204. PMC4267915

7. Zadra G, Ribeiro CF, Chetta P, Ho Y, Cacciatore S, Gao X, Syamala S, Bango C, Photopoulos C, Huang Y, Tyekucheva S, Bastos DC, Tchaicha J, Lawney B, Uo T, D’Anello L, Csibi A, Kalekar R, Larimer B, Ellis L, Butler LM, Morrissey C, McGovern K, Palombella VJ, Kutok JL, Mahmood U, Bosari S, Adams J, Peluso S, Dehm SM, Plymate SR, Loda M. Inhibition of de novo lipogenesis targets androgen receptor signaling in castration-resistant prostate cancer. Proc Natl Acad Sci U S A. 2019;116(2):631–640. PMC6329966

8. Xu L, Yin Y, Li Y, Chen X, Chang Y, Zhang H, Liu J, Beasley J, McCaw P, Zhang H, Young S, Groth J, Wang Q, Locasale JW, Gao X, Tang DG, Dong X, He Y, George D, Hu H, Huang J. A glutaminase isoform switch drives therapeutic resistance and disease progression of prostate cancer. Proc Natl Acad Sci U S A. 2021;118(13). PMC8020804

9. Massie CE, Lynch A, Ramos-Montoya A, Boren J, Stark R, Fazli L, Warren A, Scott H, Madhu B, Sharma N, Bon H, Zecchini V, Smith DM, Denicola GM, Mathews N, Osborne M, Hadfield J, Macarthur S, Adryan B, Lyons SK, Brindle KM, Griffiths J, Gleave ME, Rennie PS, Neal DE, Mills IG. The androgen receptor fuels prostate cancer by regulating central metabolism and biosynthesis. EMBO J. 2011;30(13):2719–2733. PMC3155295

10. Lin C, Blessing AM, Pulliam TL, Shi Y, Wilkenfeld SR, Han JJ, Murray MM, Pham AH, Duong K, Brun SN, Shaw RJ, Ittmann MM, Frigo DE. Inhibition of CAMKK2 impairs autophagy and castration-resistant prostate cancer via suppression of AMPK-ULK1 signaling. Oncogene. 2021;40(9):1690–1705. PMC7935762

11. Reina-Campos M, Linares JF, Duran A, Cordes T, L’Hermitte A, Badur MG, Bhangoo MS, Thorson PK, Richards A, Rooslid T, Garcia-Olmo DC, Nam-Cha SY, Salinas-Sanchez AS, Eng K, Beltran H, Scott DA, Metallo CM, Moscat J, Diaz-Meco MT. Increased Serine and One-Carbon Pathway Metabolism by PKClambda/iota Deficiency Promotes Neuroendocrine Prostate Cancer. Cancer Cell. 2019;35(3):385–400 e389. PMC6424636

12. Bader DA, Hartig SM, Putluri V, Foley C, Hamilton MP, Smith EA, Saha PK, Panigrahi A, Walker C, Zong L, Martini-Stoica H, Chen R, Rajapakshe K, Coarfa C, Sreekumar A, Mitsiades N, Bankson JA, Ittmann MM, O’Malley BW, Putluri N, McGuire SE. Mitochondrial pyruvate import is a metabolic vulnerability in androgen receptor-driven prostate cancer. Nat Metab. 2019;1(1):70–85. PMC6563330

13. Choi SYC, Ettinger SL, Lin D, Xue H, Ci X, Nabavi N, Bell RH, Mo F, Gout PW, Fleshner NE, Gleave ME, Collins CC, Wang Y. Targeting MCT4 to reduce lactic acid secretion and glycolysis for treatment of neuroendocrine prostate cancer. Cancer Med. 2018. PMC6051138

14. Rajan P, Sudbery IM, Villasevil ME, Mui E, Fleming J, Davis M, Ahmad I, Edwards J, Sansom OJ, Sims D, Ponting CP, Heger A, McMenemin RM, Pedley ID, Leung HY. Next-generation sequencing of advanced prostate cancer treated with androgen-deprivation therapy. Eur Urol. 2014;66(1):32–39. PMC4062940

15. Bishop JL, Thaper D, Vahid S, Davies A, Ketola K, Kuruma H, Jama R, Nip KM, Angeles A, Johnson F, Wyatt AW, Fazli L, Gleave ME, Lin D, Rubin MA, Collins CC, Wang Y, Beltran H, Zoubeidi A. The Master Neural Transcription Factor BRN2 Is an Androgen Receptor-Suppressed Driver of Neuroendocrine Differentiation in Prostate Cancer. Cancer Discov. 2017;7(1):54–71.

16. Divakaruni AS, Paradyse A, Ferrick DA, Murphy AN, Jastroch M. Analysis and interpretation of microplate-based oxygen consumption and pH data. Methods Enzymol. 2014;547:309–354.

17. Jones AE, Sheng L, Acevedo A, Veliova M, Shirihai OS, Stiles L, Divakaruni AS. Forces, fluxes, and fuels: tracking mitochondrial metabolism by integrating measurements of membrane potential, respiration, and metabolites. Am J Physiol Cell Physiol. 2021;320(1):C80–C91. PMC7846976

18. Desousa BR, Kim KKO, Hsieh WY, Jones AE, Swain P, Morrow DH, Ferrick DA, Shirihai OS, Neilson A, Nathanson DA, Rogers GW, Dranka BP, Murphy AN, Affourtit C, Bensinger SJ, Stiles L, Romero N, Divakaruni AS. Calculating ATP production rates from oxidative phosphorylation and glycolysis during cell activation. bioRxiv. 2022:2022.2004.2016.488523.

19. Mah CY, Nassar ZD, Swinnen JV, Butler LM. Lipogenic effects of androgen signaling in normal and malignant prostate. Asian J Urol. 2020;7(3):258–270. PMC7385522

20. Evans KW, Yuca E, Scott SS, Zhao M, Paez Arango N, Cruz Pico CX, Saridogan T, Shariati M, Class CA, Bristow CA, Vellano CP, Zheng X, Gonzalez-Angulo AM, Su X, Tapia C, Chen K, Akcakanat A, Lim B, Tripathy D, Yap TA, Francesco MED, Draetta GF, Jones P, Heffernan TP, Marszalek JR, Meric-Bernstam F. Oxidative Phosphorylation Is a Metabolic Vulnerability in Chemotherapy-Resistant Triple-Negative Breast Cancer. Cancer Res. 2021;81(21):5572–5581. PMC8563442

21. Palanisamy N, Yang J, Shepherd PDA, Li-Ning-Tapia EM, Labanca E, Manyam GC, Ravoori MK, Kundra V, Araujo JC, Efstathiou E, Pisters LL, Wan X, Wang X, Vazquez ES, Aparicio AM, Carskadon SL, Tomlins SA, Kunju LP, Chinnaiyan AM, Broom BM, Logothetis CJ, Troncoso P, Navone NM. The MD Anderson Prostate Cancer Patient-derived Xenograft Series (MDA PCa PDX) Captures the Molecular Landscape of Prostate Cancer and Facilitates Marker-driven Therapy Development. Clin Cancer Res. 2020;26(18):4933–4946. PMC7501166

22. Quigley DA, Dang HX, Zhao SG, Lloyd P, Aggarwal R, Alumkal JJ, Foye A, Kothari V, Perry MD, Bailey AM, Playdle D, Barnard TJ, Zhang L, Zhang J, Youngren JF, Cieslik MP, Parolia A, Beer TM, Thomas G, Chi KN, Gleave M, Lack NA, Zoubeidi A, Reiter RE, Rettig MB, Witte O, Ryan CJ, Fong L, Kim W, Friedlander T, Chou J, Li H, Das R, Li H, Moussavi-Baygi R, Goodarzi H, Gilbert LA, Lara PN, Jr., Evans CP, Goldstein TC, Stuart JM, Tomlins SA, Spratt DE, Cheetham RK, Cheng DT, Farh K, Gehring JS, Hakenberg J, Liao A, Febbo PG, Shon J, Sickler B, Batzoglou S, Knudsen KE, He HH, Huang J, Wyatt AW, Dehm SM, Ashworth A, Chinnaiyan AM, Maher CA, Small EJ, Feng FY. Genomic Hallmarks and Structural Variation in Metastatic Prostate Cancer. Cell. 2018;175(3):889.

23. Davies A, Nouruzi S, Ganguli D, Namekawa T, Thaper D, Linder S, Karaoglanoglu F, Omur ME, Kim S, Kobelev M, Kumar S, Sivak O, Bostock C, Bishop J, Hoogstraat M, Talal A, Stelloo S, van der Poel H, Bergman AM, Ahmed M, Fazli L, Huang H, Tilley W, Goodrich D, Feng FY, Gleave M, He HH, Hach F, Zwart W, Beltran H, Selth L, Zoubeidi A. An androgen receptor switch underlies lineage infidelity in treatment-resistant prostate cancer. Nat Cell Biol. 2021;23(9):1023–1034. PMC9012003

24. Stine ZE, Walton ZE, Altman BJ, Hsieh AL, Dang CV. MYC, Metabolism, and Cancer. Cancer Discov. 2015;5(10):1024–1039. PMC4592441

25. Akamatsu S, Wyatt AW, Lin D, Lysakowski S, Zhang F, Kim S, Tse C, Wang K, Mo F, Haegert A, Brahmbhatt S, Bell R, Adomat H, Kawai Y, Xue H, Dong X, Fazli L, Tsai H, Lotan TL, Kossai M, Mosquera JM, Rubin MA, Beltran H, Zoubeidi A, Wang Y, Gleave ME, Collins CC. The Placental Gene PEG10 Promotes Progression of Neuroendocrine Prostate Cancer. Cell Rep. 2015;12(6):922–936.

26. Gomes LC, Di Benedetto G, Scorrano L. During autophagy mitochondria elongate, are spared from degradation and sustain cell viability. Nat Cell Biol. 2011;13(5):589–598. PMC3088644

27. Petcherski A, Trudeau KM, Wolf DM, Segawa M, Lee J, Taddeo EP, Deeney JT, Liesa M. Elamipretide Promotes Mitophagosome Formation and Prevents Its Reduction Induced by Nutrient Excess in INS1 beta-cells. J Mol Biol. 2018;430(24):4823–4833. PMC6290358

28. Chan DC. Mitochondrial Dynamics and Its Involvement in Disease. Annu Rev Pathol. 2020;15:235–259.

29. Mishra P, Chan DC. Metabolic regulation of mitochondrial dynamics. J Cell Biol.2016;212(4):379–387. PMC4754720

30. Lee YG, Nam Y, Shin KJ, Yoon S, Park WS, Joung JY, Seo JK, Jang J, Lee S, Nam D, Caino MC, Suh PG, Chan Chae Y. Androgen-induced expression of DRP1 regulates mitochondrial metabolic reprogramming in prostate cancer. Cancer Lett. 2020;471:72–87.

31. Rambold AS, Kostelecky B, Elia N, Lippincott-Schwartz J. Tubular network formation protects mitochondria from autophagosomal degradation during nutrient starvation. Proc Natl Acad Sci U S A. 2011;108(25):10190–10195. PMC3121813

32. Li J, Huang Q, Long X, Guo X, Sun X, Jin X, Li Z, Ren T, Yuan P, Huang X, Zhang H, Xing J. Mitochondrial elongation-mediated glucose metabolism reprogramming is essential for tumour cell survival during energy stress. Oncogene. 2017;36(34):4901–4912.

33. Molina JR, Sun Y, Protopopova M, Gera S, Bandi M, Bristow C, McAfoos T, Morlacchi P, Ackroyd J, Agip AA, Al-Atrash G, Asara J, Bardenhagen J, Carrillo CC, Carroll C, Chang E, Ciurea S, Cross JB, Czako B, Deem A, Daver N, de Groot JF, Dong JW, Feng N, Gao G, Gay J, Do MG, Greer J, Giuliani V, Han J, Han L, Henry VK, Hirst J, Huang S, Jiang Y, Kang Z, Khor T, Konoplev S, Lin YH, Liu G, Lodi A, Lofton T, Ma H, Mahendra M, Matre P, Mullinax R, Peoples M, Petrocchi A, Rodriguez-Canale J, Serreli R, Shi T, Smith M, Tabe Y, Theroff J, Tiziani S, Xu Q, Zhang Q, Muller F, DePinho RA, Toniatti C, Draetta GF, Heffernan TP, Konopleva M, Jones P, Di Francesco ME, Marszalek JR. An inhibitor of oxidative phosphorylation exploits cancer vulnerability. Nat Med. 2018;24(7):1036–1046.

34. He L, Wondisford FE. Metformin action: concentrations matter. Cell Metab. 2015;21(2):159–162.

35. Mukhopadhyay S, Goswami D, Adiseshaiah PP, Burgan W, Yi M, Guerin TM, Kozlov SV, Nissley DV, McCormick F. Undermining Glutaminolysis Bolsters Chemotherapy While NRF2 Promotes Chemoresistance in KRAS-Driven Pancreatic Cancers. Cancer Res. 2020;80(8):1630–1643. PMC7185043

36. Zhou W, Yao Y, Scott AJ, Wilder-Romans K, Dresser JJ, Werner CK, Sun H, Pratt D, Sajjakulnukit P, Zhao SG, Davis M, Nelson BS, Halbrook CJ, Zhang L, Gatto F, Umemura Y, Walker AK, Kachman M, Sarkaria JN, Xiong J, Morgan MA, Rehemtualla A, Castro MG, Lowenstein P, Chandrasekaran S, Lawrence TS, Lyssiotis CA, Wahl DR. Purine metabolism regulates DNA repair and therapy resistance in glioblastoma. Nat Commun. 2020;11(1):3811. PMC7393131

37. Baek ML, Lee J, Pendleton KE, Berner MJ, Goff EB, Tan L, Martinez SA, Wang T, Meyer MD, Lim B, Barrish JP, Porter W, Lorenzi PL, Echeverria GV. Mitochondrial structure and function adaptation in residual triple negative breast cancer cells surviving chemotherapy treatment. bioRxiv. 2022:2022.2002.2025.481996.

38. Kong B, Tsuyoshi H, Orisaka M, Shieh DB, Yoshida Y, Tsang BK. Mitochondrial dynamics regulating chemoresistance in gynecological cancers. Ann N Y Acad Sci. 2015;1350:1–16.

39. Courtois S, de Luxan-Delgado B, Penin-Peyta L, Royo-Garcia A, Parejo-Alonso B, Jagust P, Alcala S, Rubiolo JA, Sanchez L, Sainz B, Jr., Heeschen C, Sancho P. Inhibition of Mitochondrial Dynamics Preferentially Targets Pancreatic Cancer Cells with Enhanced Tumorigenic and Invasive Potential. Cancers (Basel). 2021;13(4). PMC7914708

40. Yu M, Nguyen ND, Huang Y, Lin D, Fujimoto TN, Molkentine JM, Deorukhkar A, Kang Y, San Lucas FA, Fernandes CJ, Koay EJ, Gupta S, Ying H, Koong AC, Herman JM, Fleming JB, Maitra A, Taniguchi CM. Mitochondrial fusion exploits a therapeutic vulnerability of pancreatic cancer. JCI Insight. 2019;5. PMC6777817

41. Hagenbuchner J, Kuznetsov AV, Obexer P, Ausserlechner MJ. BIRC5/Survivin enhances aerobic glycolysis and drug resistance by altered regulation of the mitochondrial fusion/fission machinery. Oncogene. 2013;32(40):4748–4757.

42. Ku SY, Rosario S, Wang Y, Mu P, Seshadri M, Goodrich ZW, Goodrich MM, Labbe DP, Gomez EC, Wang J, Long HW, Xu B, Brown M, Loda M, Sawyers CL, Ellis L, Goodrich DW. Rb1 and Trp53 cooperate to suppress prostate cancer lineage plasticity, metastasis, and antiandrogen resistance. Science. 2017;355(6320):78–83. PMC5367887

43. Mu P, Zhang Z, Benelli M, Karthaus WR, Hoover E, Chen CC, Wongvipat J, Ku SY, Gao D, Cao Z, Shah N, Adams EJ, Abida W, Watson PA, Prandi D, Huang CH, de Stanchina E, Lowe SW, Ellis L, Beltran H, Rubin MA, Goodrich DW, Demichelis F, Sawyers CL. SOX2 promotes lineage plasticity and antiandrogen resistance in TP53- and RB1-deficient prostate cancer. Science. 2017;355(6320):84–88. PMC5247742

44. Guo H, Ci X, Ahmed M, Hua JT, Soares F, Lin D, Puca L, Vosoughi A, Xue H, Li E, Su P, Chen S, Nguyen T, Liang Y, Zhang Y, Xu X, Xu J, Sheahan AV, Ba-Alawi W, Zhang S, Mahamud O, Vellanki RN, Gleave M, Bristow RG, Haibe-Kains B, Poirier JT, Rudin CM, Tsao MS, Wouters BG, Fazli L, Feng FY, Ellis L, van der Kwast T, Berlin A, Koritzinsky M, Boutros PC, Zoubeidi A, Beltran H, Wang Y, He HH. ONECUT2 is a driver of neuroendocrine prostate cancer. Nat Commun. 2019;10(1):278. PMC6336817

45. Aggarwal R, Huang J, Alumkal JJ, Zhang L, Feng FY, Thomas GV, Weinstein AS, Friedl V, Zhang C, Witte ON, Lloyd P, Gleave M, Evans CP, Youngren J, Beer TM, Rettig M, Wong CK, True L, Foye A, Playdle D, Ryan CJ, Lara P, Chi KN, Uzunangelov V, Sokolov A, Newton Y, Beltran H, Demichelis F, Rubin MA, Stuart JM, Small EJ. Clinical and Genomic Characterization of Treatment-Emergent Small-Cell Neuroendocrine Prostate Cancer: A Multi-institutional Prospective Study. J Clin Oncol. 2018;36(24):2492–2503. PMC6366813

46. Beltran H, Prandi D, Mosquera JM, Benelli M, Puca L, Cyrta J, Marotz C, Giannopoulou E, Chakravarthi BV, Varambally S, Tomlins SA, Nanus DM, Tagawa ST, Van Allen EM, Elemento O, Sboner A, Garraway LA, Rubin MA, Demichelis F. Divergent clonal evolution of castration-resistant neuroendocrine prostate cancer. Nat Med. 2016;22(3):298–305. PMC4777652

47. Demir U, Koehler A, Schneider R, Schweiger S, Klocker H. Metformin anti-tumor effect via disruption of the MID1 translational regulator complex and AR downregulation in prostate cancer cells. BMC Cancer. 2014;14:52. PMC3929757

48. Ben Sahra I, Laurent K, Loubat A, Giorgetti-Peraldi S, Colosetti P, Auberger P, Tanti JF, Le Marchand-Brustel Y, Bost F. The antidiabetic drug metformin exerts an antitumoral effect in vitro and in vivo through a decrease of cyclin D1 level. Oncogene. 2008;27(25):3576–3586.

49. Colquhoun AJ, Venier NA, Vandersluis AD, Besla R, Sugar LM, Kiss A, Fleshner NE, Pollak M, Klotz LH, Venkateswaran V. Metformin enhances the antiproliferative and apoptotic effect of bicalutamide in prostate cancer. Prostate Cancer Prostatic Dis. 2012;15(4):346–352.

50. Nobes JP, Langley SE, Klopper T, Russell-Jones D, Laing RW. A prospective, randomized pilot study evaluating the effects of metformin and lifestyle intervention on patients with prostate cancer receiving androgen deprivation therapy. BJU Int. 2012;109(10):1495–1502.

51. Ahn HK, Lee YH, Koo KC. Current Status and Application of Metformin for Prostate Cancer: A Comprehensive Review. Int J Mol Sci. 2020;21(22). PMC7698147

52. Hess C, Unger M, Madea B, Stratmann B, Tschoepe D. Range of therapeutic metformin concentrations in clinical blood samples and comparison to a forensic case with death due to lactic acidosis. Forensic Sci Int. 2018;286:106–112.

