## Supplementary material for "Androgen receptor inhibition induces metabolic reprogramming and increased reliance on oxidative mitochondrial metabolism in prostate cancer": Materials and Methods

#### 13C isotope tracing

Naïve and LTenza 16D cells were plated in 6-well dishes at 225,000 and 350,000 cells/well respectively. After 24 hours, cells were washed and cultured in a base RPMI supplemented with 10 mM uniformly labeled 13C6-glucose (Cambridge Isotope Laboratories), 10% (v/v) FBS, 2 mM glutamine, 100 units/mL penicillin, and 100 µg/mL streptomycin.

24 hours after the addition of U-13$C_6$-glucose, cells were harvested and extracted for GC/MS analysis using established methods[1]. Briefly, cell plates were placed on ice and quickly washed with ice-cold 0.9% (w/v) NaCl. Cells were immediately treated with 500 µL of ice-cold MeOH and 200 µL water containing 1 µg of the internal standard norvaline. Cells were then scraped and placed in 1.5 mL centrifuge tubes kept on ice. Next, 500 µL of chloroform was added, after which samples were vortexed for 1 min and then spun at 10,000*g* for 5 min at 4C. The aqueous layer was transferred to a GC/MS sample vial and dried overnight using a refrigerated CentriVap.

Once dry, samples were resuspended in 20 µL of 2% (w/v) methoxyamine in pyridine and incubated at 37C for 45 minutes. This was followed by addition of 20 µL of MTBSTFA + 1% TBDMSCl (Ntert-Butyldimethylsilyl-N-methyltrifluoroacetamide with 1% tertButyldimethylchlorosilane), mixing, and incubation for an additional 45 minutes at 37C. Samples were run as previously described[1], and analyzed using Agilent MassHunter software. Stable isotope tracing data was corrected for natural abundance of heavy isotopes with FluxFix software using a reference set of unlabeled metabolite standards[2].

#### Animal work

All animal work was performed using IACUC approved protocols under the supervision of veterinarians from the Division of Laboratory Animal Medicine at UCLA. 7 million 16D cells were implanted subcutaneously with 100 µl Matrigel (Corning) into NSG mice to form primary tumors. Primary tumors were harvested, minced, and re-implanted (20 - 80 mg of minced tumor tissue with 100ul Matrigel per mouse) into NSG mice. 16D tumor-bearing mice were treated by oral gavage with 10 mg/kg/day of enzalutamide in the vehicle (1% carboxymethyl cellulose, 0.5% Tween 80, and 5% dimethylsulfoxide) or the vehicle only, with a two-days-on/one-day-off schedule. Tumors were collected after 10 days of treatment and prepared for histology, protein extraction, and metabolite extraction. 180-30 PDX tumors were maintained by serial implantation of 20 - 80 mg of minced tumor tissue. Treatment with vehicle or enzalutamide was initiated one week after implantation and performed using the approach described above. Tumors were collected after 7 days of treatment and prepared for protein extraction and *ex vivo* organoid culture.

#### Apoptosis analysis

Cells were seeded at 40 percent confluence and cultured in 6-well dishes for 48 hours prior to apoptosis analysis. No media changes were performed to preserve all material. Cell culture media and wash media were collected and pooled with quenched trypsincontaining media containing cells and apoptosis analysis was performed using an apoptosis detection kit (BioLegend, 640922) according to the provided protocol. Flow cytometry was performed to quantify the percentage of annexin $V^+$ cells. In experiments using transduced lines, analysis was restricted to the transduced cells which were identified via flow cytometry by analyzing RFP florescence.

**Bioenergetic assays**

Oxygen consumption and extracellular acidification rates were measured using an Agilent Seahorse XF96 or $XF^e$96 Analyzer. Briefly, 16D prostate cancer cells were plated at 40,000 cells/well in XF96 plates for 24 hours. At the time of experiment, tissue culture growth medium was replaced with assay medium consisting of unbuffered DMEM (Sigma, 5030) supplemented with 10mM glucose, 2mM pyruvate, 2mM glutamine, and 5mM HEPES. Respiration was measured at baseline and in response to acute treatment with 2 μM oligomycin, FCCP (two sequential pulses of 500nM), and 0.2μM rotenone with 1 μM antimycin A. All respiratory parameters were calculated as previously described in[3]. Where indicated, cells were treated with 5μM UK5099, 5μM etomoxir, or 1μM CB-839 15 minutes prior to recording the initial measurements.

Rates of ATP produced from oxidative phosphorylation and glycolysis were calculated as previously described[4]. Mitochondrial ATP production rates were determined by stoichiometric conversion of the ATP-linked respiration rate, and glycolytic ATP production rates were measured by correcting rates of extracellular acidification for the scaling factor of the microplate sensor coverage and confounding respiratory acidification. Where indicated, cells were treated with 2 μM oligomycin 15 minutes prior to recording the initial measurements.

**Cell cycle analysis**

Cells were seeded at 30 percent confluence and cultured in 6-well dishes for 72 hours prior to cell cycle analysis. Media changes were performed 48 hours after plating. After 72 hours of culture, cell cycle analysis was performed using a 5-ethynyl-2'-deoxyuridine-based (EdU) kit (ThermoFisher, C10635) according to the specified protocol. EdU labeling was performed for 2 hours in LNCaP and 16D cells, and for 5 hours in 180-30 PDX organoids. For experiments that contained small molecule inhibitors, fresh inhibitor(s) were adding during each media change. PDX 180-30 organoids were dissociated after EdU labeling prior to fixation for cell cycle analysis. In select experiments, the 1ug/ml Hoechst 33342 DNA stain (ThermoFisher, 62249) was added prior to flow cytometry analysis to identify G2 and M phase cells. Flow cytometry analysis identified the percentage of EdU-positive and/or Hoechst-positive cells. For experiments with transduced lines, analysis was restricted to the RFP-positive transduced cells.

**Cell lines, lentiviral transductions, and cloning of knockdown vectors**

Cell lines were routinely tested for mycoplasma and authentication by short tandem repeat analysis (Laragen). Tissue culture plates were coated with 0.01% (v/v) Poly-L-Lysine (Sigma, P4832) diluted 1/20 in distilled water and washed with PBS to enhance cell attachment. 16D and LNCaP cells were cultured in RPMI base media (Gibco) + 10% FBS (v/v) + 100 units/mL penicillin, and 100 μg/mL streptomycin. Enzalutamide treatment was performed by adding 10uM enzalutamide (Selleck Chemicals, S1250) every 48 hours. For LNCaP castration experiments, LNCaP cells were grown in RPMI base media + 10% CSS (v/v) (Sigma, F6765) + 100 units/mL penicillin, and 100 µg/mL streptomycin +/- 0.5nM DHT (Sigma) and fresh media was provided every 48 hours.

For lentiviral transductions, cells were seeded at 30-50% confluence. Transductions were performed 24-48 hours after seeding with 8ug/ml Polybrene (Fisher, NC0663391). Flow cytometry-based cell sorting was performed at least 72 hours after transduction to isolate color-tagged transduced cells.

MYC virus was produced using a plasmid vector FU-MYC-CRW[5]. Plasmid vectors for shHK2, and DRP1S616E were obtained from VectorBuilder. The target sequence for HK2 is TGACGACAGCATCATTGTTAA. shScramble (FU-shScr-CRW) and shRB1-shTP53 (FU-shTP35-shRB1-CRW) vectors were cloned as follows. First, the pBSPacI shuttle vector was made by inserting an adaptor oligonucleotide AG220: 5'-AATTCTTTAATTAAAG-3' at the EcoRI site of pBluescript II KS(+) (Stratagene). The pPass1 shuttle was cloned as follows. Annealed oligonucleotides AG232: 5'-CCTTAATTAAGCGATCGCACTGGGTACCTGGGCC-3' and AG233: 5'-CAGGTACCCAGTGCGATCGCTTAATTAAGGGTAC-3' were inserted between KpnI and ApaI sites of pBluescript II KS(+). Then, annealed oligonucleotides AG234: 5'-CTTAATTAAACTGGGGAGCTCCGC-3' and AG235: 5'-GGAGCTCCCCAGTTTAATTAAGAGCT-3' were inserted between SacI and SacII sites. This creates a PacI-AsiSI-[Multiple cloning sites]-PacI cassette. Annealed oligonucleotides AG218: 5'-GACGATGATTAATTAA-3', and AG220 (above) were ligated with KflI-EcoRI fragment of shp53 pLO1 pure (Addgene) and inserted into PacI site in pBSPacI (pBSPacI-shTP53). PacI fragment of pBSPacI-shTP53 was then inserted into the AsiSI site in pPassI (pPass1-shTP53). The PacI-PacI fragment of FU-shRB1-AR-CGW was blunted and digested with HindIII creating a HindIII-blunt fragment of H1-shRB1 cassette. This cassette was inserted between HindIII and EcoRV sites in pPass1-shTP53 (pPass1-shTP53-shRB1). PacI fragment of pPass1-shTP53-shRB1 was then inserted into the PacI site of FU-CRW (FU-shTP53-shRB1-CRW). The U6-Scramble cassette was made by ligating annealed oligonucleotides AG227: 5'-CACCGAATTCTTCCATAGAGCTCGTCAAGAGCGAGCTCTATGGAAGAATTC-3' and AG228: 5'-AAAAGAATTCTTCCATAGAGCTCGCTCTTGACGAGCTCTATGGAAGAATTC-3' in pENTR/U6 vector (Invitrogen). Then, the BamHI-XbaI fragment was purified and inserted between BamHI and XbaI sites in pBSPacI (pBSPacI-shScr21A). The PacI fragment from pBSPacI-shScr21A was inserted at the PacI site in FU-CRW (FU-shScr-CRW). Concentrated viral aliquots were produced either by VectorBuilder or UCLA Integrated Molecular Technologies Core.

**ChIP sequencing**
AR ChIP-seq bigwig files were generated using deeptools program suite[6]. AR binding profiles of LNCaP-Ctrl, LNCaP-R1881[7], 16D[8] samples at genomic loci (namely LDHA, HK2 and DNM1L) were analyzed by visualizing AR ChIP-seq bigwig tracks using IGV[9].

**DNA methylation analysis**
Bisulfite sequencing was carried out as described previously[10]. In brief, genomic DNA was bisulfite converted using the EZ DNA methylation kit (Zymo Research, Orange, CA, USA) and amplified using primers specific to the promoter of HK2 (F-5′-AGTTGAGTTTTAGTGATTTTGTGGT -3′, R-5′-AACTTACCTTCTACACTTAATCATAATTAA -3). All PCR reactions were carried out in 40 µl volume containing 20 ng of bisulfite converted DNA, 1 × Platinum Taqbuffer (Invitrogen), 1.5 U Platinum Taq (Life Technologies), 250 µM each dNTPs, 1.5 mM MgCl2, 0.25 µg/µl bovine serum albumin, 2 µl dimethyl sulfoxide, 400 nM forward primer, and 400 nM reverse primer. Cycling conditions were 95C for 3 min, 36 cycles of 95C for 30 s, 55C for 30 s, and 72C for 30 s, followed by a 7 min extension step at 72C. PCR products were gel purified after electrophoresis on a 2% agarose gel. Amplicons were sequenced to an average coverage of 27,380x using established amplicon sequencing protocols (Azenta). Raw bisulfite amplicon sequencing fastq reads were first trimmed using Trim Galore version 0.6.6 and then aligned to UCSC hg19 reference genome using Bismark version 0.23.0[11]. Bismark was further used to deduplicate the alignments and extract methylation call files which report the percentage of methylated cytosines for each CpG dinucleotide.

**Immunohistochemistry**
16D tumor tissue from vehicle- and 10-day enzalutamide-treated mice was fixed with 4% paraformaldehyde in PBS for 6-8 hours and processed for paraffin sections. Tumor samples were color inked and formalin-fixed paraffin embedded. 3- to 4-µm thick sections were placed on charged slides for immunohistochemical staining that was carried out on Dako's Automated AS48Link Autostainer in SPORE Pathology Core laboratory at UCLA. Positive and negative control slides were pretreated with Heat Induced Epitope Retrieval (HIER) in Dako PT Link using the Envision FLEX Target Retrieval solution at low pH (6.0), and incubated at 97C for 15 minutes. Primary Rabbit anti-Human HK2 monoclonal antibody (Cell Signaling, 28675), clone C64G5, was added at a 1:600 dilution and incubated for 60 minutes at room temperature. Sections were then incubated for 5 minutes with the EnVision Flex+ Rabbit linker (Agilent, SM805) prior to a 5-minute treatment with the Polymer Flex/HRP (Agilent, SM802) reagent. Negative control slides received the Flex Rabbit Negative Control Immunoglobulin fraction (Agilent, IR600) instead of primary antibody. Slides were developed in Envision Flex DAB+Chromogen and counterstained with Mayer's hematoxylin.

***In vivo* metabolomics**
After tumor dissection, a maximum of 30mg of tissue was weighed, snap frozen, and stored at -80C until metabolite extraction. To extract metabolites, weighed tumor tissue was added to a bead tube (Fisher) containing 1ml 80% methanol plus 10mM potassium trifluoromethanesulfonate (TMSO) internal standard on ice. Samples were homogenized for 1 minute at max speed on a bead homogenizer (Fisher). Bead tubes were spun at 17000g at 4C for 10 minutes. The supernatant was transferred to an Eppendorf tube and spun at 17000g at 4C for 10 minutes. A volume of extraction equivalent to 3mg of tumor tissue was transferred to an ABC vial (Fisher). All volumes were normalized to 500ul with 80% methanol containing TMSO internal standard. 80% MeOH was evaporated from the

ABC vials using the EZ-2Elite evaporator (Genevac) and samples were stored at -80°C until analysis.

Dried metabolites were reconstituted in 100 µL of a 50% acetonitrile (ACN) 50% dH20 solution. Samples were vortexed and spun down for 10 min at 17,000g. 70 µL of the supernatant was then transferred to HPLC glass vials. 10 µL of these metabolite solutions were injected per analysis. Samples were run on a Vanquish (Thermo Scientific) UHPLC system with mobile phase A (20mM ammonium carbonate, pH 9.7) and mobile phase B (100% ACN) at a flow rate of 150 µL/min on a SeQuant ZIC-pHILIC Polymeric column (2.1 × 150 mm 5 µm, EMD Millipore) at 35°C. Separation was achieved with a linear gradient from 20% A to 80% A in 20 min followed by a linear gradient from 80% A to 20% A from 20 min to 20.5 min. 20% A was then held from 20.5 min to 28 min. The UHPLC was coupled to a Q-Exactive (Thermo Scientific) mass analyzer running in polarity switching mode with spray-voltage=3.2kV, sheath-gas=40, aux-gas=15, sweep-gas=1, aux-gas-temp=350°C, and capillary-temp=275°C. For both polarities mass scan settings were kept at full-scan-range=(70-1000), ms1-resolution=70,000, max-injection-time=250ms, and AGC-target=1E6. MS2 data was also collected from the top three most abundant singly-charged ions in each scan with normalized-collision-energy=35. Each of the resulting ".RAW" files was then centroided and converted into two ".mzXML" files (one for positive scans and one for negative scans) using msconvert from ProteoWizard[12]. These ".mzXML" files were imported into the MZmine 2 software package[13]. Ion chromatograms were generated from MS1 spectra via the built-in Automated Data Analysis Pipeline (ADAP) chromatogram module[14] and peaks were detected via the ADAP wavelets algorithm. Peaks were aligned across all samples via the Random sample consensus aligner module, gap-filled, and assigned identities using an exact mass MS1(+/-15ppm) and retention time RT (+/-0.5min) search of our in-house MS1-RT database. Peak boundaries and identifications were then further refined by manual curation. Peaks were quantified by area under the curve integration and exported as CSV files. If stable isotope tracing was used in the experiment, the peak areas were additionally processed via the R package AccuCor[15] to correct for natural isotope abundance. Peak areas for each sample were normalized by the measured area of the internal standard trifluoromethanesulfonate (present in the extraction buffer) and by the number of cells present in the extracted well.

***In vitro* metabolomic profiling**

Cells were seeded at 30 percent confluence and cultured in 6-well dishes for 72 hours prior to metabolite extractions. Media was aspirated and cells were washed with cold 150mM ammonium acetate pH 7.3. Metabolite extractions were performed by adding 500ul of cold 80% methanol containing 2nM Norvaline (Sigma) as an internal standard per well. Cells were removed using a cell scraper before transferring cell suspensions to 1.5ml Eppendorf tubes. Samples were vortexed for 30 seconds and spun at 4C for 5 minutes at maximum speed to pellet the insoluble fraction before 420ul of the soluble fraction was transferred to ABC vials (Fisher). 80% MeOH was evaporated from the ABC vials using the EZ-2Elite evaporator (Genevac) and samples were stored at -80°C until analysis.

Dried metabolites were resuspended in 50% ACN:water and 1/10$^{th}$ was loaded onto a Luna 3um NH2 100A (150 × 2.0 mm) column (Phenomenex). The chromatographic separation was performed on a Vanquish Flex (Thermo Scientific) with mobile phases A (5 mM NH4AcO pH 9.9) and B (ACN) and a flow rate of 200 µl/min. A linear gradient from 15% A to 95% A over 18 min was followed by 9 min isocratic flow at 95% A and reequilibration to 15% A. Metabolites were detection with a Thermo Scientific Q Exactive mass spectrometer run with polarity switching (+3.5 kV/− 3.5 kV) in full scan mode with an m/z range of 70-975 and 70.000 resolution. TraceFinder 4.1 (Thermo Scientific) was used to quantify the targeted metabolites by area under the curve using expected retention time and accurate mass measurements (< 5 ppm).

Normalization was performed by resuspending the insoluble fraction in 300ul of lysis solution (0.1M NaCl, 20mM Tris-HCl, 0.1% SDS, 5mM EDTA in distilled water) and proceeding with DNA measurement. Samples were syringed with a 25G needle to reduce viscosity and 50ul of each sample was transferred to a 96-well black wall clear bottom tissue culture plate (Corning). 50ul lysis solution was added to one well for a blank reading. 100ul of 5ug/ml Hoechst 33342 (Invitrogen) in distilled water was added to each well and 96-well plates were incubated for 30 minutes in the dark at 37C before measurement of DNA-based florescence using a Tecan Infinite M1000 plate reader with 355nm excitation and 465nm emission. The blank reading was subtracted from each absorbance value to calculate relative cell amount.

**Organoid culture**

Using a razor blade, individual tumors were mechanically dissociated in dissociation media comprised of RPMI-1640 containing 10% (v/v) fetal bovine serum, 100 units/mL penicillin, and 100 µg/mL streptomycin, 1mg/mL collagenase type I, 1mg/ml dispase, 0.1mg/mL deoxyribonuclease, and 10uM of the p160ROCK inhibitor Y-27632 dihydrochloride (Tocris Bioscience). When chunks were no longer visible, the samples were incubated at 37C on a nutating platform for 45 minutes in 10mL of dissociation media. After centrifugation at 800g for 5 min, the pellet was washed with 1x phosphate buffered saline. The cell pellet was resuspended in human organoid media[16] and passed through a 100um cell strainer. Growth factor reduced Matrigel (Corning) was added to the cell suspension at a final concentration of 75% before plating into rings in 24-well plates. After Matrigel rings solidified at 37C for 1 hour, 500ul human organoid media was added to each well. Each vehicle- and enzalutamide-treated sample was cultured +/- 30nM IACS-010759 (ChemieTek) for 72 hours.

**RNA sequencing**

RNA was extracted from samples using the RNeasy Mini Kit (QIAGEN). Library preparation was performed using the KAPA Stranded mRNA-Seq Kit (Roche). The workflow consists of mRNA enrichment, cDNA generation, and end repair to generate blunt ends, A-tailing, adaptor ligation, and PCR amplification. Different adaptors were used for multiplexing samples in one lane. The Illumina HiSeq 3000 was used to perform sequencing for 1x50 run.

#### Visualization of mitochondria

Cells were cultured in μ-Slide Well (Ibidi) and fixed with 4% PSA/PBS for 2 minutes. After washing with PBS, cells were stained with anti-TUFM (Atlas Antibodies, AMAb90966) followed by Alexa Fluor 488-conjugated anti-mouse IgG (H+L) (Invitrogen, A11001) and 4',6-diamidino-2-phenylindole (DAPI) (Sigma, D8417). Signals were visualized using Zeiss LSM 880 confocal scanning microscope with Airyscan with 100x oil immersion objectives.

#### Western blot

Cells were lysed in RIPA buffer (50mM Tris-HCL pH8.0, 150nM NaCl, 1% NP-40, 0.5% Sodium Deoxycholate, 0.1% SDS) containing a phosphatase inhibitor cocktail (Halt, 78428) and a protease inhibitor cocktail (Millipore Sigma, 11697498001). Sonication was performed with a sonic dismembrator (Fisher, FB120) to improve membranous and nuclear protein yield. For extraction of protein lysate from tumor samples, tumors were minced with a razor blade prior to transfer to pre-filled bead mill tubes (Fisher, 15-340-153) and resuspension in the lysis solution described above. Homogenization was performed for 2 minutes at max intensity using a Bead Mill 4 homogenizer (Fisher, 15-340-164). Samples were run on NuPAGE 4%-12% Bis-Tris Gels (Fisher, NP0335) and protein was transferred to PVDF transfer membranes (Fisher, IPV00010). Total protein was visualized using the SYPRO RUBY protein blot stain (Fisher, S11791) and membranes were blocked in PBS +.1% Tween-20 (Fisher, BP337-500) + 5% milk (Fisher, BC9121673). Proteins were probed with primary antibodies followed by chromophore-conjugated anti-mouse (Invitrogen, A21235) or anti-rabbit secondary antibodies (Invitrogen, A21244) or HRP-conjugated anti-mouse (Thermo, 31430) or anti-rabbit secondary antibodies (Thermo, 31463) and detected via florescence or HRP chemiluminescence respectively. Primary antibodies used were beta-Actin (Invitrogen, MA1-140), Androgen Receptor (Cell Signaling, 5153S), Hexokinase II (Cell Signaling, 28675), DRP1 (Cell Signaling, 5391S), Phospho-DRP1 (Ser616) (Cell Signaling, 3455S), Anti-LDH-A (MilliporeSigma, MABC150), Recombinant-Anti-c-MYC (Abcam, ab32072), NSE (Proteintech, 66150-1-Ig), and PSA/KLK3 (Cell Signaling, 5877).

### QUANTIFICATION AND STATISTICAL ANALYSIS

#### Metabolomics analysis

For projection plots, principal component analysis (PCA) was performed with the scikit-learn, NumPy, pandas, and Matplotlib libraries in Python. Feature selection was done based on shared features between differing datasets. Count/abundance matrices were sorted along their respective feature-axis to ensure features were listed in the same order. After performing z-score scaling, the coordinates from the *in vitro* samples were merged onto a PCA plot with the values from *in vivo* samples. 95% confidence ellipses were generated from PCA-transformed coordinates using a script from Matplotlib (https://github.com/Nick-Nunley/PCA-for-AR-induced-metabolic-reprogramming-in-CRPCa). Heatmaps were generated by plotting row z-scores in GraphPad Prism Version 7. To generate the average z-score plot, an *in vivo* enza-enriched metabolite signature was defined. Row z-scores of *in vivo* enza-enriched metabolites were calculated from *the in vitro* metabolomics dataset. Row z-scores from three technical replicates from a representative experiment (n=3) were averaged and represented on a dot plot. MSEA was generated using Metaboanalyst 5.0[17] (https://www.metaboanalyst.ca/MetaboAnalyst/home.xhtml).

**Mitochondrial content and morphology**
Mitochondrial elongation was expressed as aspect ratio (long axis/short axis ratio) and eccentricity, calculated as the ratio of the distance between the foci of an ellipse and its major axis length. Branching was expressed as form factor $((4\pi(\text{area}))/\text{Perimeter}^2)$. Mitochondrial parameters were determined from mitochondrial TUFM staining. Image analysis was performed using ImageJ v1.53c and CellProfiler v2.0[18]. For mitochondrial volume quantification, z-stack images were processed with Imaris software (Oxford Instruments) to identify TUFM-positive regions and calculate TUFM-positive volume.

**RNA sequencing analysis**
KEGG pathway analysis was performed using DAVID Bioinformatics[19,20]. GSEA was performed as described previously using GSEA_4.0.3 software[21,22]. Projection plots were generated as described for metabolomics analysis. After performing z-score scaling, the coordinates from the *in vitro* 16D enzalutamide time-course RNA-sequencing data were merged onto a PCA plot with the values from the *Rajan et al* dataset. 95% confidence ellipses were generated as described for metabolomics analysis. Average z-score plots and heatmaps were generated as described for metabolomics analysis.

In the *Quigley et al* dataset, there were 63 enzalutamide-naive and 36 enzalutamide-resistant patients whose tumor underwent RNA-seq[23]. Alignment to hg38-decoy reference was performed using STAR aligner (version 2.5.0b) with per-gene counts quantification on the basis of Illumina RNA-seq alignment app Version 1.1.0[24].

**Western blot quantification**
Western blots were quantified using ImageJ software. Background values were subtracted from the mean gray value for each band. Each band was normalized to its respective loading control.

### DATA AND CODE AVAILABILITY
The data discussed in this publication have been deposited in NCBI’s Gene Expression Omnibus and are accessible through GEO Series accession numbers GSE202885, GSE202755, and GSE202897.

### MATERIALS AND METHODS REFERENCES

1. Cordes T, Metallo CM. Quantifying Intermediary Metabolism and Lipogenesis in Cultured Mammalian Cells Using Stable Isotope Tracing and Mass Spectrometry. *Methods Mol Biol.* 2019;1978:219-241.
2. Trefely S, Ashwell P, Snyder NW. FluxFix: automatic isotopologue normalization for metabolic tracer analysis. *BMC Bioinformatics.* 2016;17(1):485. PMC5123363

3. Divakaruni AS, Paradyse A, Ferrick DA, Murphy AN, Jastroch M. Analysis and interpretation of microplate-based oxygen consumption and pH data. *Methods Enzymol.* 2014;547:309-354.
4. Desousa BR, Kim KKO, Hsieh WY, Jones AE, Swain P, Morrow DH, Ferrick DA, Shirihai OS, Neilson A, Nathanson DA, Rogers GW, Dranka BP, Murphy AN, Affourtit C, Bensinger SJ, Stiles L, Romero N, Divakaruni AS. Calculating ATP production rates from oxidative phosphorylation and glycolysis during cell activation. *bioRxiv.* 2022:2022.2004.2016.488523.
5. Stoyanova T, Cooper AR, Drake JM, Liu X, Armstrong AJ, Pienta KJ, Zhang H, Kohn DB, Huang J, Witte ON, Goldstein AS. Prostate cancer originating in basal cells progresses to adenocarcinoma propagated by luminal-like cells. *Proc Natl Acad Sci U S A.* 2013;110(50):20111-20116. PMC3864278
6. Ramirez F, Ryan DP, Gruning B, Bhardwaj V, Kilpert F, Richter AS, Heyne S, Dundar F, Manke T. deepTools2: a next generation web server for deep-sequencing data analysis. *Nucleic Acids Res.* 2016;44(W1):W160-165. PMC4987876
7. Zhang A, Zhao JC, Kim J, Fong KW, Yang YA, Chakravarti D, Mo YY, Yu J. LncRNA HOTAIR Enhances the Androgen-Receptor-Mediated Transcriptional Program and Drives Castration-Resistant Prostate Cancer. *Cell Rep.* 2015;13(1):209-221. PMC4757469
8. Davies A, Nouruzi S, Ganguli D, Namekawa T, Thaper D, Linder S, Karaoglanoglu F, Omur ME, Kim S, Kobelev M, Kumar S, Sivak O, Bostock C, Bishop J, Hoogstraat M, Talal A, Stelloo S, van der Poel H, Bergman AM, Ahmed M, Fazli L, Huang H, Tilley W, Goodrich D, Feng FY, Gleave M, He HH, Hach F, Zwart W, Beltran H, Selth L, Zoubeidi A. An androgen receptor switch underlies lineage infidelity in treatment-resistant prostate cancer. *Nat Cell Biol.* 2021;23(9):1023-1034. PMC9012003
9. Robinson JT, Thorvaldsdottir H, Winckler W, Guttman M, Lander ES, Getz G, Mesirov JP. Integrative genomics viewer. *Nat Biotechnol.* 2011;29(1):24-26. PMC3346182
10. Yegnasubramanian S, Lin X, Haffner MC, DeMarzo AM, Nelson WG. Combination of methylated-DNA precipitation and methylation-sensitive restriction enzymes (COMPARE-MS) for the rapid, sensitive and quantitative detection of DNA methylation. *Nucleic Acids Res.* 2006;34(3):e19. PMC1363782
11. Krueger F, Andrews SR. Bismark: a flexible aligner and methylation caller for Bisulfite-Seq applications. *Bioinformatics.* 2011;27(11):1571-1572. PMC3102221
12. Chambers MC, Maclean B, Burke R, Amodei D, Ruderman DL, Neumann S, Gatto L, Fischer B, Pratt B, Egertson J, Hoff K, Kessner D, Tasman N, Shulman N, Frewen B, Baker TA, Brusniak MY, Paulse C, Creasy D, Flashner L, Kani K, Moulding C, Seymour SL, Nuwaysir LM, Lefebvre B, Kuhlmann F, Roark J, Rainer P, Detlev S, Hemenway T, Huhmer A, Langridge J, Connolly B, Chadick T, Holly K, Eckels J, Deutsch EW, Moritz RL, Katz JE, Agus DB, MacCoss M, Tabb DL, Mallick P. A cross-platform toolkit for mass spectrometry and proteomics. *Nat Biotechnol.* 2012;30(10):918-920. PMC3471674

13. Pluskal T, Castillo S, Villar-Briones A, Oresic M. MZmine 2: modular framework for processing, visualizing, and analyzing mass spectrometry-based molecular profile data. *BMC Bioinformatics.* 2010;11:395. PMC2918584
14. Myers OD, Sumner SJ, Li S, Barnes S, Du X. One Step Forward for Reducing False Positive and False Negative Compound Identifications from Mass Spectrometry Metabolomics Data: New Algorithms for Constructing Extracted Ion Chromatograms and Detecting Chromatographic Peaks. *Anal Chem.* 2017;89(17):8696-8703.
15. Su X, Lu W, Rabinowitz JD. Metabolite Spectral Accuracy on Orbitraps. *Anal Chem.* 2017;89(11):5940-5948. PMC5748891
16. Drost J, Karthaus WR, Gao D, Driehuis E, Sawyers CL, Chen Y, Clevers H. Organoid culture systems for prostate epithelial and cancer tissue. *Nat Protoc.* 2016;11(2):347-358. PMC4793718
17. Pang Z, Chong J, Zhou G, de Lima Morais DA, Chang L, Barrette M, Gauthier C, Jacques PE, Li S, Xia J. MetaboAnalyst 5.0: narrowing the gap between raw spectra and functional insights. *Nucleic Acids Res.* 2021;49(W1):W388-W396. PMC8265181
18. Kamentsky L, Jones TR, Fraser A, Bray MA, Logan DJ, Madden KL, Ljosa V, Rueden C, Eliceiri KW, Carpenter AE. Improved structure, function and compatibility for CellProfiler: modular high-throughput image analysis software. *Bioinformatics.* 2011;27(8):1179-1180. PMC3072555
19. Jacobs PP, Geysens S, Vervecken W, Contreras R, Callewaert N. Engineering complex-type N-glycosylation in Pichia pastoris using GlycoSwitch technology. *Nat Protoc.* 2009;4(1):58-70.
20. Huang da W, Sherman BT, Lempicki RA. Bioinformatics enrichment tools: paths toward the comprehensive functional analysis of large gene lists. *Nucleic Acids Res.* 2009;37(1):1-13. PMC2615629
21. Subramanian A, Tamayo P, Mootha VK, Mukherjee S, Ebert BL, Gillette MA, Paulovich A, Pomeroy SL, Golub TR, Lander ES, Mesirov JP. Gene set enrichment analysis: a knowledge-based approach for interpreting genome-wide expression profiles. *Proc Natl Acad Sci U S A.* 2005;102(43):15545-15550. PMC1239896
22. Mootha VK, Lindgren CM, Eriksson KF, Subramanian A, Sihag S, Lehar J, Puigserver P, Carlsson E, Ridderstrale M, Laurila E, Houstis N, Daly MJ, Patterson N, Mesirov JP, Golub TR, Tamayo P, Spiegelman B, Lander ES, Hirschhorn JN, Altshuler D, Groop LC. PGC-1alpha-responsive genes involved in oxidative phosphorylation are coordinately downregulated in human diabetes. *Nat Genet.* 2003;34(3):267-273.
23. Quigley DA, Dang HX, Zhao SG, Lloyd P, Aggarwal R, Alumkal JJ, Foye A, Kothari V, Perry MD, Bailey AM, Playdle D, Barnard TJ, Zhang L, Zhang J, Youngren JF, Cieslik MP, Parolia A, Beer TM, Thomas G, Chi KN, Gleave M, Lack NA, Zoubeidi A, Reiter RE, Rettig MB, Witte O, Ryan CJ, Fong L, Kim W, Friedlander T, Chou J, Li H, Das R, Li H, Moussavi-Baygi R, Goodarzi H, Gilbert LA, Lara PN, Jr., Evans CP, Goldstein TC, Stuart JM, Tomlins SA, Spratt DE, Cheetham RK, Cheng DT, Farh K, Gehring JS, Hakenberg J, Liao A, Febbo PG, Shon J, Sickler B, Batzoglou S, Knudsen KE, He HH, Huang J, Wyatt AW, Dehm SM, Ashworth A, Chinnaiyan

AM, Maher CA, Small EJ, Feng FY. Genomic Hallmarks and Structural Variation in Metastatic Prostate Cancer. *Cell.* 2018;174(3):758-769 e759. PMC6425931

24. Dobin A, Davis CA, Schlesinger F, Drenkow J, Zaleski C, Jha S, Batut P, Chaisson M, Gingeras TR. STAR: ultrafast universal RNA-seq aligner. *Bioinformatics.* 2013;29(1):15-21. PMC3530905
