## Supplementary Figures for "Androgen receptor inhibition induces metabolic reprogramming and increased reliance on oxidative mitochondrial metabolism in prostate cancer"

### Supplementary Figure 1 (related to Figure 1)

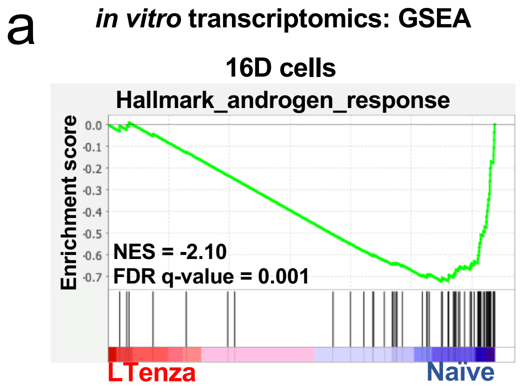


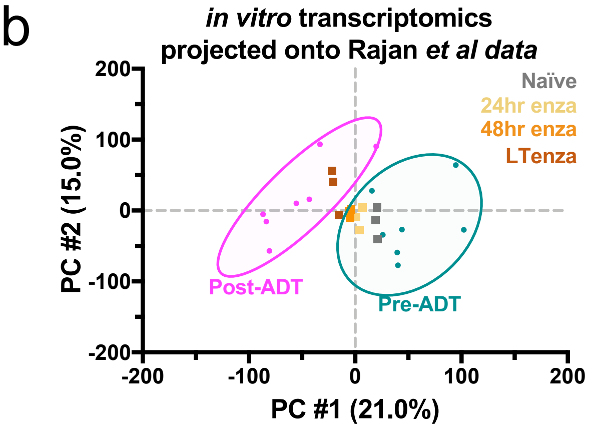


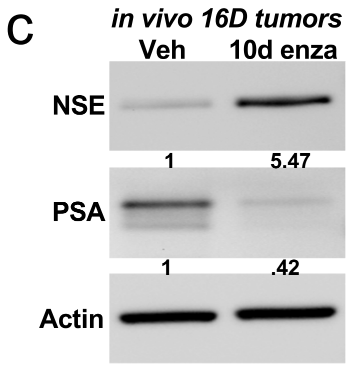


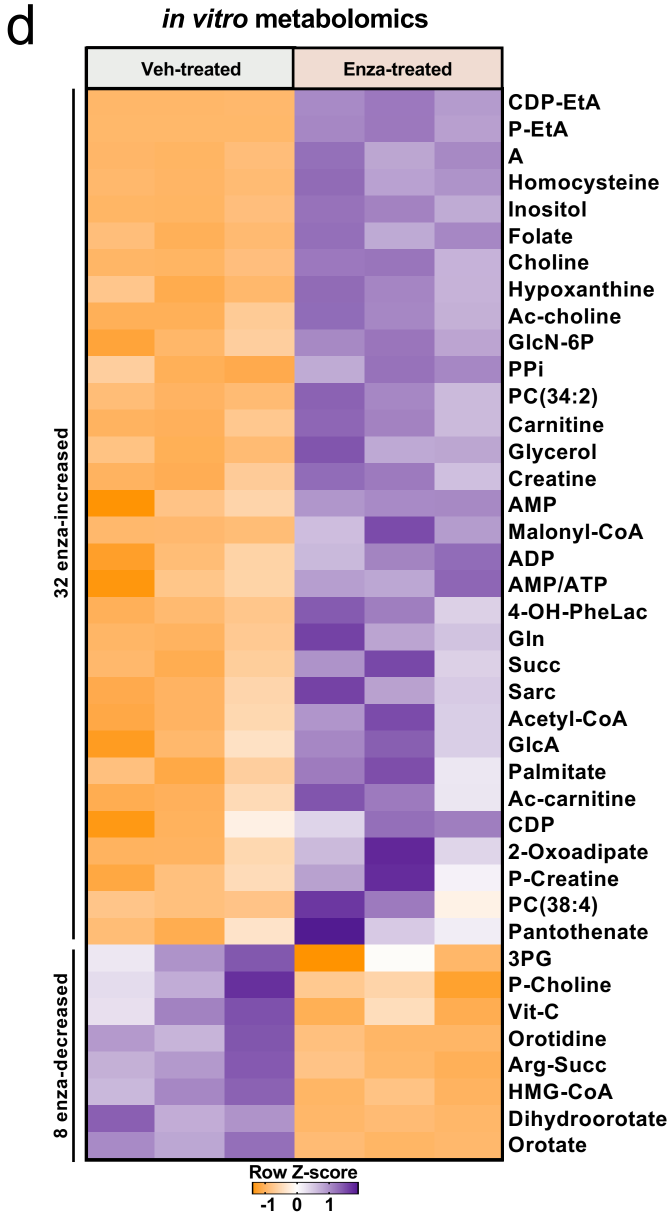

**Supplementary Figure 1. Validation of *in vitro* 16D model, *in vivo* enzalutamide delivery and heatmap from *in vitro* metabolomics.** (a) Gene Set Enrichment Analysis (GSEA) of Hallmark_androgen_response genes in naïve and LTenza 16D cells showing normalized enrichment score (NES) and false discovery rate (FDR). (b) Naïve, 24hr enzalutamide-treated (enza), 48hr enza, and LTenza 16D transcriptomics data projected onto principle component analysis (PCA) plot of pre-androgen deprivation therapy (pre-ADT) and post-ADT samples from *Rajan et al* data. (c) Western blot analysis of NSE, PSA, and Actin (loading control) in lysates from vehicle-treated (Veh) and 10-day (10d) enzalutamide-treated subcutaneous 16D tumors. (d) Heatmap of differentially abundant metabolites (fold change ≥ 1.25, FDR < 0.05) in LTenza 16D cells (Enza-treated) compared to naïve (Veh-treated) 16D cells.

### Supplementary Figure 2 (related to Figure 3)

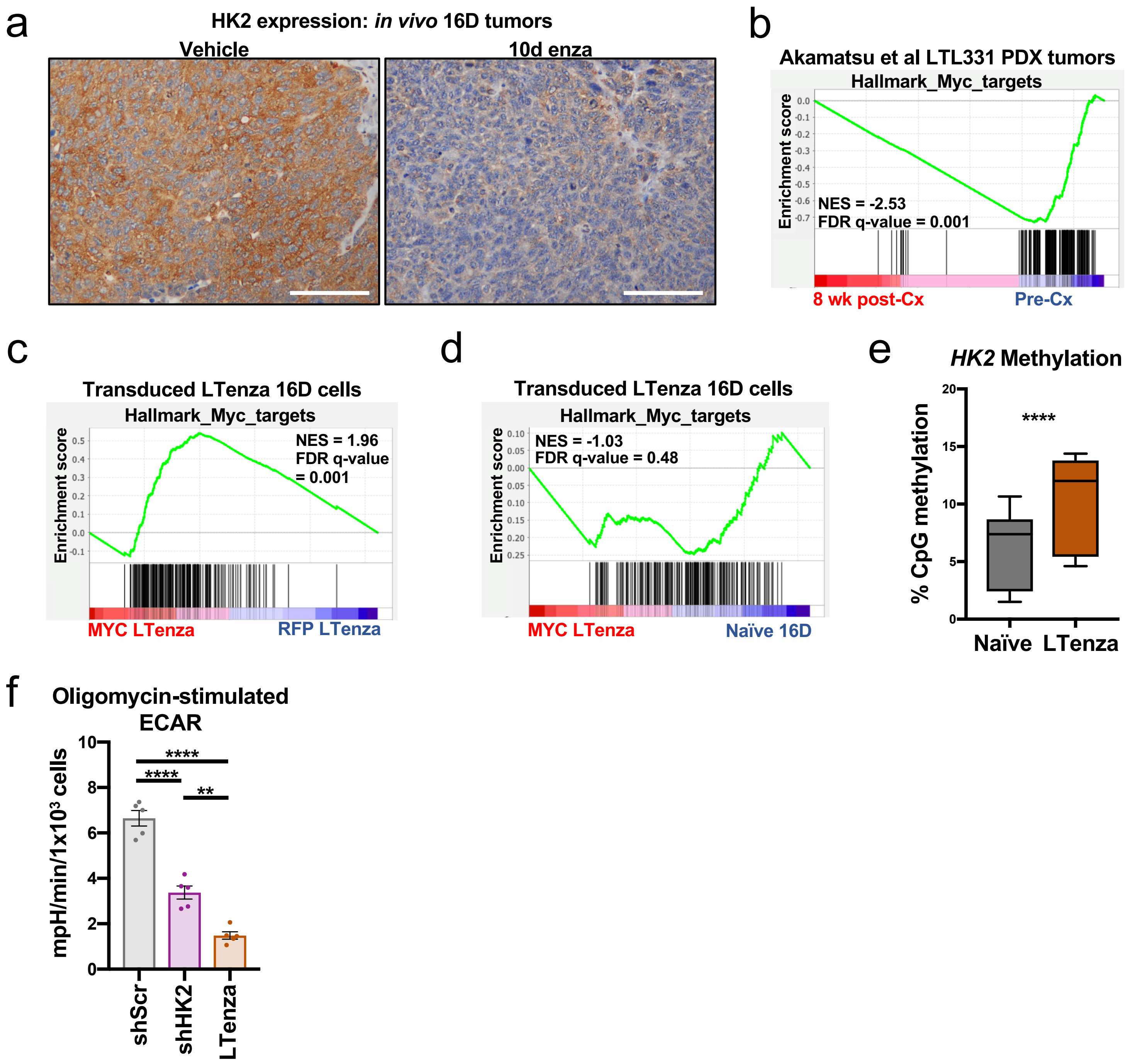

**Supplementary Figure 2. Reduced MYC activity and reduced glycolysis through HK2 downregulation in anti-androgen treated cells.** (a) 3, 3'-diaminobenzidine staining indicates HK2 expression in representative tissue sections from vehicle- and 10-day enzalutamide-treated (10d enza) 16D tumors. Scale bars, 100$\mu$m. (b) GSEA of Hallmark_Myc_targets in pre-castration (pre-Cx) and 8-week post-castration (8 wk post-Cx) LTL331 tumor samples from the *Akamatsu et al* dataset. (c) GSEA of Hallmark_Myc_targets in RFP-transduced LTenza (RFP LTenza) and MYC-transduced LTenza (MYC LTenza) 16D cells. (d) GSEA of Hallmark_Myc_targets in naïve 16D and MYC LTenza 16D cells. (e) Mean percentage of methylated CpGs within the *HK2* locus of naïve and LTenza 16D cells. (f) Oligomycin-stimulated Extracellular Acidification Rate (ECAR) of shScr-transduced naïve, shHK2-transduced naïve, and LTenza 16D cells. Data represent the mean +/- SEM of 5 technical replicates from a representative experiment (n=2). Significance was evaluated (b-d) using normalized enrichment scores (NES) and false discovery rates (FDR). P-values were calculated from an unpaired t-test with Welch's correction (e and f). **p < 0.01, ****p < 0.0001.

Supplementary Figure 3 (related to Figure 4)

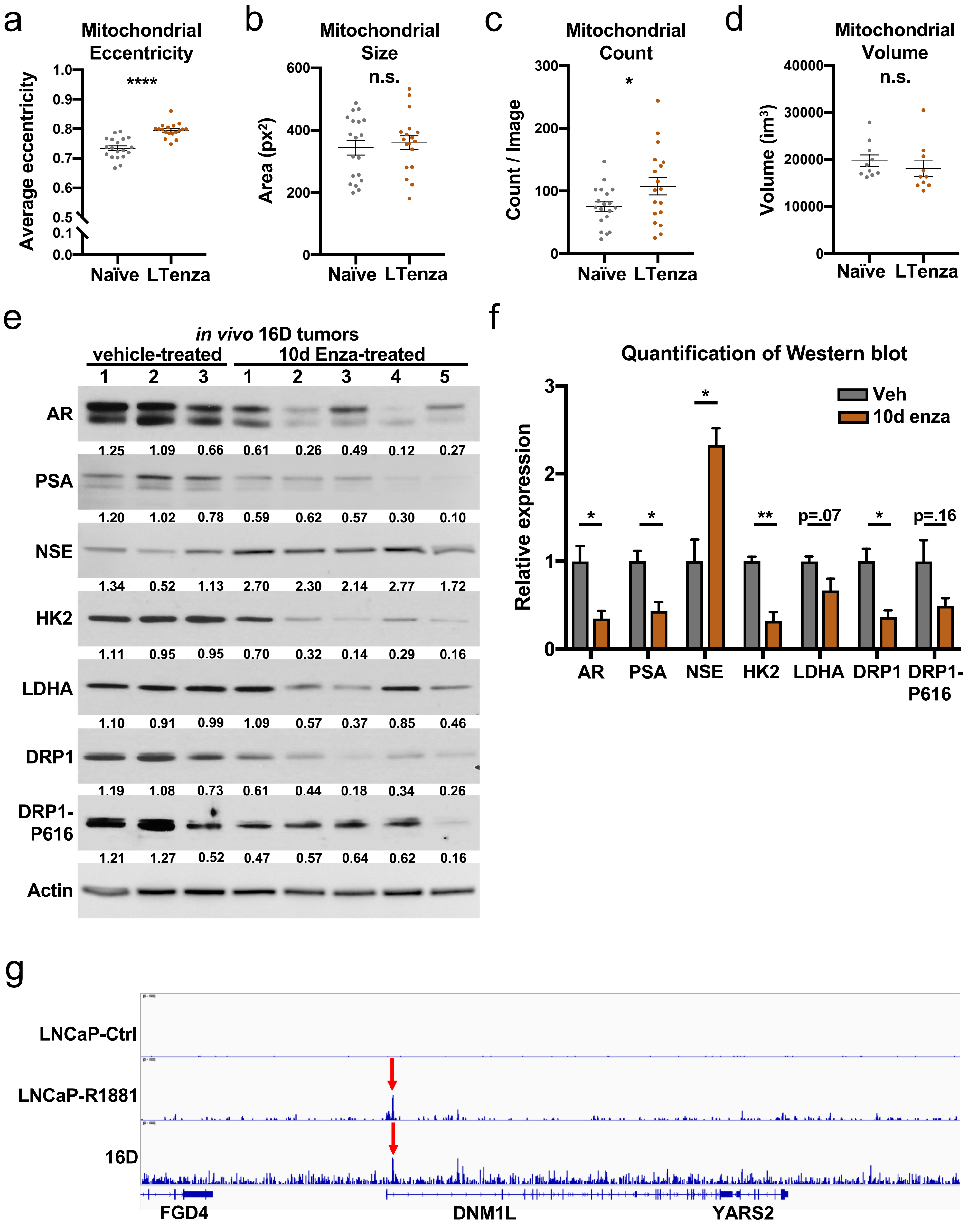

**Supplementary Figure 3. Quantification of mitochondrial parameters in naïve and LTenza 16D cells and DRP1 expression in vehicle- and enzalutamide-treated 16D tumors.** (a-c) Quantification of mitochondrial eccentricity (a), mitochondrial size (b), and mitochondrial count (c) from TUFM stains from 20 images of naïve or LTenza 16D cells. Data represent the mean +/- SEM. (d) Quantification of the mitochondrial volume of naïve and LTenza 16D cells from 3-dimensional reconstruction of 10 z-stack images per treatment group. Data represent the mean +/- SEM. (e and f) Western blot indicating AR, PSA, NSE, HK2, LDHA, DRP1, DRP1 phosphorylation at S616 (DRP1-P616), and Actin (control) expression in lysates from 3 vehicle-treated and 5 10-day enzalutamide-treated (10d Enza-treated) 16D tumors (e) and associated quantification (f). Data represent the mean +/- SEM. (g) AR binding of LNCaP-Ctrl, LNCaP-R1881 and 16D at the DNM1L genomic locus was analyzed by visualizing AR ChIP-seq bigwig tracks. Red arrow indicates sharp peak called by macs to demonstrate binding of AR. P-values were calculated from an unpaired t-test with Welch's correction. $^{*}p < 0.05$, $^{**}p < 0.01$, $^{****}p < 0.0001$, n.s. = not significant, $p \geq 0.05$.

Supplementary Figure 4 (related to Figure 5)

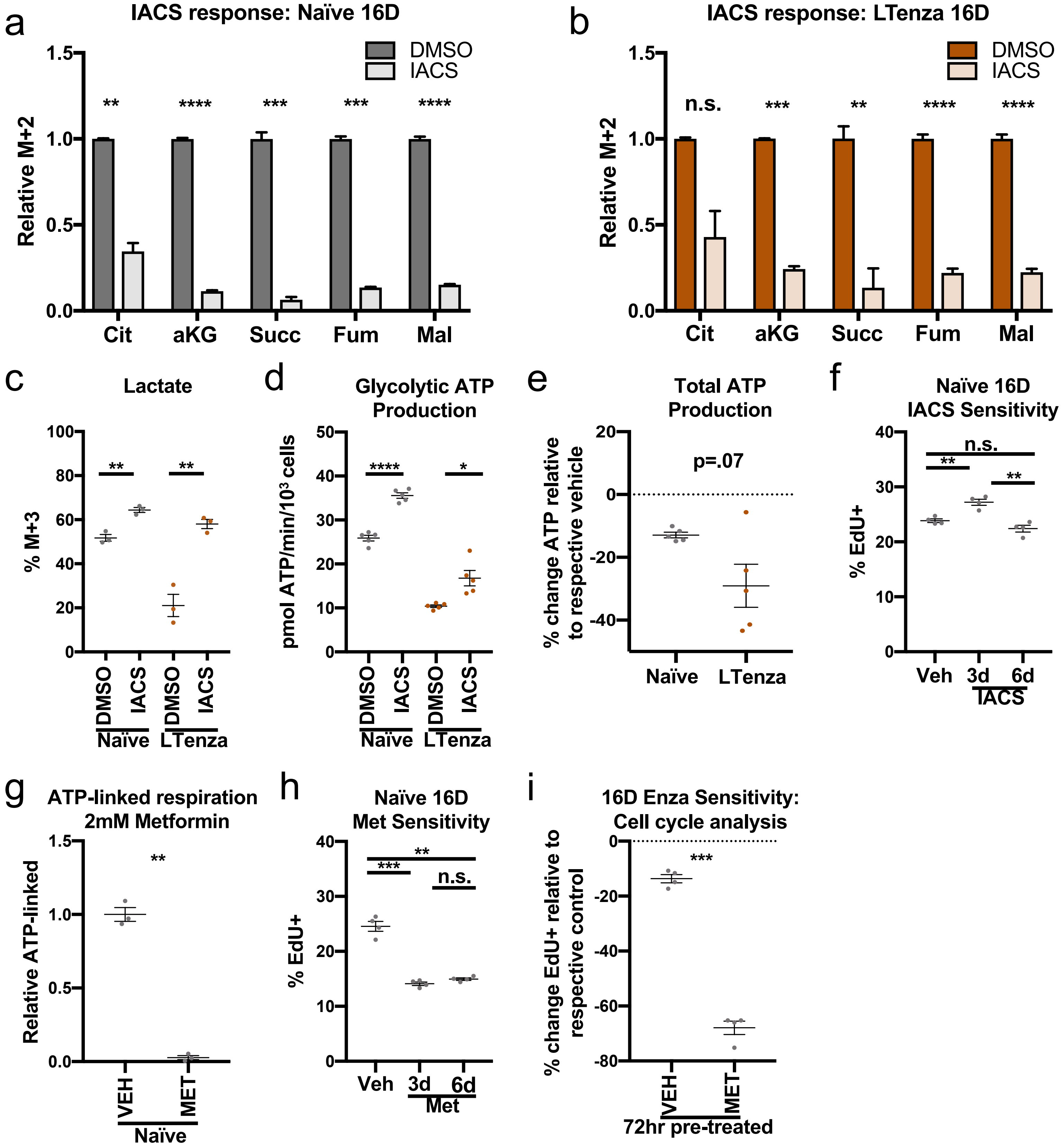

**Supplementary Figure 4. Validation of complex I inhibitors and sensitivity of enzalutamide-treated cells to complex I inhibition.** (a and b) Relative M+2-labeling of citrate (cit), alpha-ketoglutarate (aKG), succinate (succ), fumarate (fum), and malate after 24hr U-13$C_6$-glucose tracer analysis of DMSO- and 30nM IACS-010759-treated naïve (a) or LTenza 16D cells (b). Data represent the mean +/- SEM of 3 technical replicates. (c) Percentage of M+3-labeled lactate in naïve and LTenza 16D cells treated with DMSO or 30nM IACS for 24 hours. Data represent the mean +/- SEM of 3 technical replicates. (d) Glycolytic ATP production of naïve and LTenza 16D cells treated with DMSO or 30nM IACS for 24 hours. Data represent the mean +/- SEM of 5 technical replicates. (e) Relative sensitivity of the total ATP production of naïve and LTenza 16D cells to 30nM IACS. Data represent the mean +/- SEM of 5 technical replicates. (f) Cell cycle analysis measuring the proliferation (% $EdU^+$) of vehicle-, 3-day (3d), and 6d 30nM IACS-treated naïve 16D cells. Data represent the mean +/- SEM of 4 technical replicates from a representative experiment (n=3). (g) ATP-linked respiration of naïve 16D cells treated with DMSO or 2mM metformin (Met) for 24 hrs. Data represent the mean +/- SEM of 3 technical replicates. (h) Cell cycle analysis measuring the percentage of $EdU^+$ cells of vehicle-, 3d, and 6d 2mM Met-treated naïve 16D cells. Data represent the mean +/- SEM of 4 technical replicates. (i) Cell cycle analysis to quantify the relative sensitivity of DMSO- and 72hr 2mM Met-treated naïve 16D cells to enzalutamide. Data represent the mean +/- SEM of 4 technical replicates. P-values were calculated from an unpaired t-test with Welch's correction. $^*p < 0.05$, $^{**}p < 0.01$, $^{***}p < 0.001$, $^{****}p < 0.0001$, n.s. = not significant, $p \geq 0.05$.

Supplementary Figure 5 (related to Figure 5)

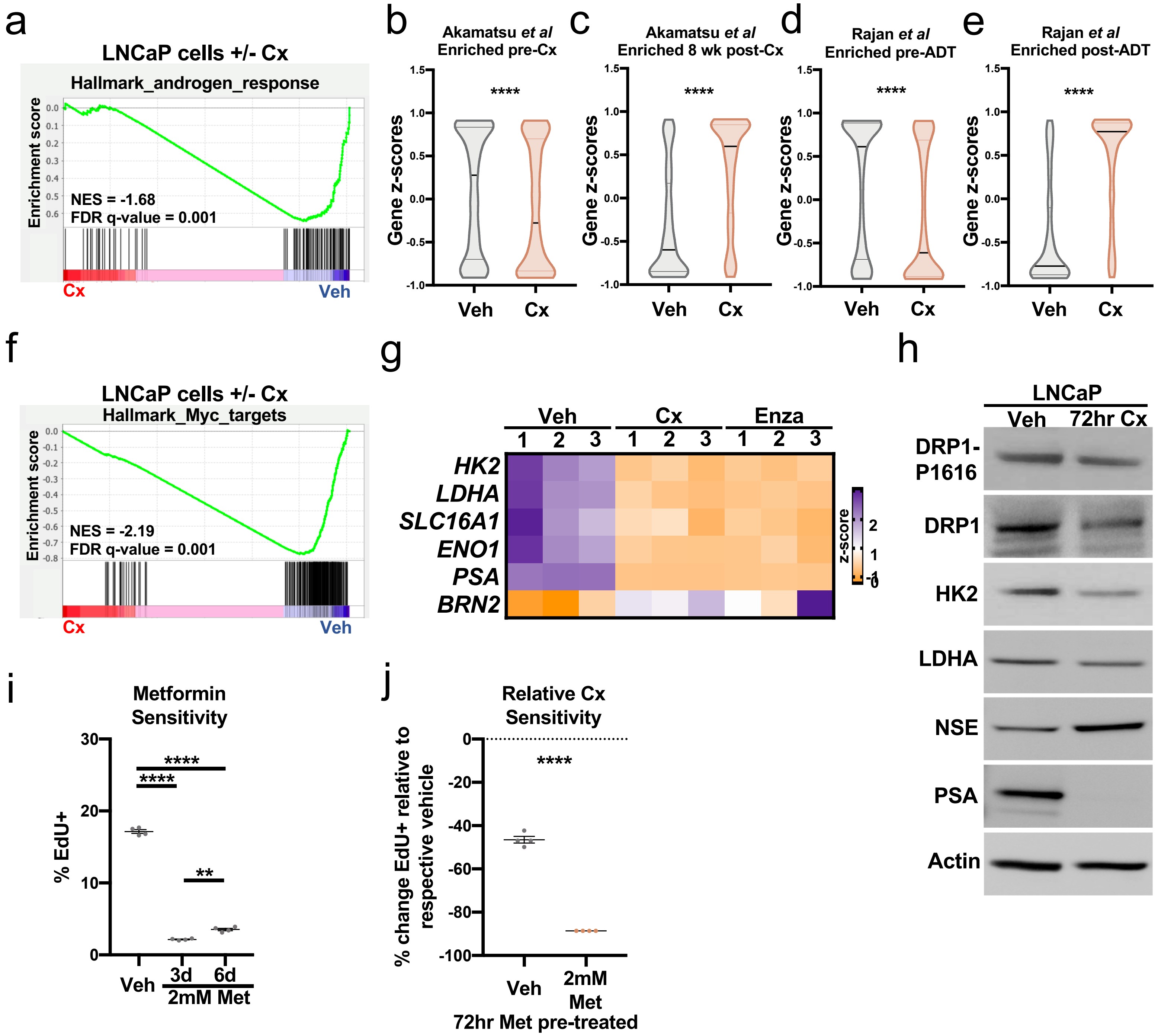

**Supplementary Figure 5. AR inhibition-induced metabolic hallmarks are observed in LNCaP cells after castration.** (a) GSEA of Hallmark_androgen_response genes in control (veh) and 72-hour castrated (Cx) LNCaP cells showing normalized enrichment score (NES) and false discovery rate (FDR). (b and c) Violin plots indicating gene z-scores of *Akamatsu et al* genes enriched pre-Cx (fold change ≥ 3, row mean > 1, 2710 genes) (b), or 8 weeks post-Cx (fold change ≥ 3, row mean > 1, 3371 genes) (c) in veh and 72-hour Cx LNCaP cells. Data represent mean +/- SEM. (d and e) Violin plots indicating gene z-scores of *Rajan et al* genes enriched pre-ADT (fold change ≥ 2, FDR < 0.2, 911 genes) (d) or post-ADT (fold change ≥ 2, FDR < 0.2, 1023 genes) (e) in veh and 72-hour Cx LNCaP cells. Data represent mean +/- SEM. (f) GSEA of Hallmark_Myc_targets genes in veh and 72-hour Cx LNCaP cells showing NES and FDR. (g) Heatmap showing the mRNA expression of select glycolytic genes from RNA sequencing of 3 technical replicates of veh, 72-hour Cx, and 72-hour enzalutamide-treated (enza) LNCaP cells. (h) Western blot detecting DRP1 phosphorylation at S616 (DRP1-P616), DRP1, HK2, LDHA, NSE, PSA, and Actin (control) in veh and 72-hour Cx LNCaP lysates. (i) Cell cycle analysis measuring the % $EdU^{+}$ cells of veh, 3-day (3d), and 6d 2mM metformin-treated (Met) LNCaP cells. Data represent the mean +/- SEM of 4 technical replicates. (j) Cell cycle analysis to quantify the relative sensitivity of veh and 72hr Met LNCaP cells to castration. Data represent the mean +/- SEM of 4 technical replicates. P-values were calculated from an unpaired t-test with Welch's correction. **p < 0.01, ****p < 0.0001.

Supplementary Figure 6

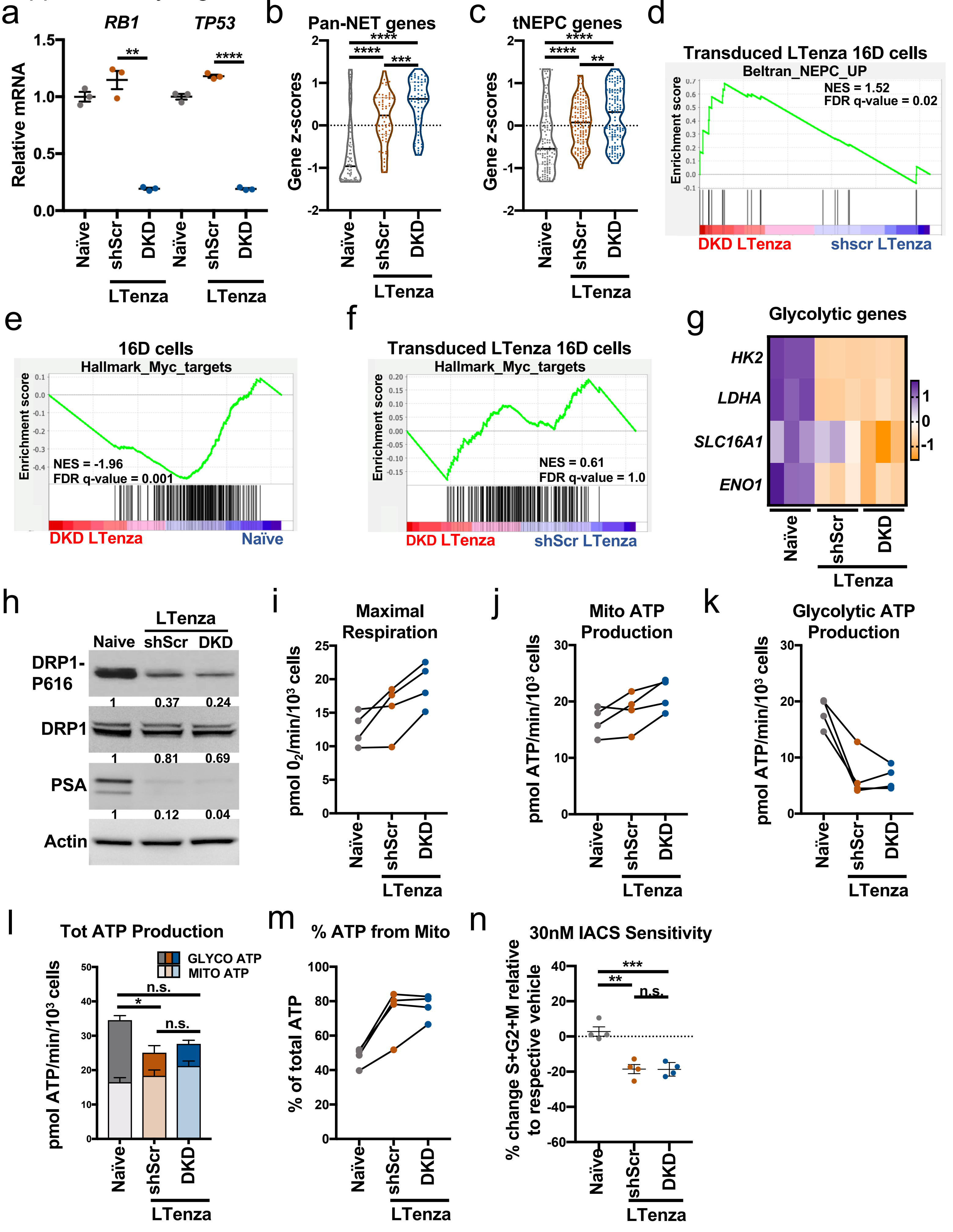

**Supplementary Figure 6. AR inhibition-induced metabolic hallmarks are retained in enzalutamide-treated 16D cells after knockdown of *RB1* and *TP53*.** (a) mRNA expression of *RB1* and *TP53* in naïve, shScr-transduced (shScr) LTenza, and shRB1_shTP53-transduced (DKD) LTenza 16D cells. Data represent the mean +/- SEM of 3 technical replicates. (b and c) Violin plots indicating gene z-scores of Pan-neuroendocrine tumor (Pan-NET) associated genes defined by *Guo et al* (b) and treatment-emergent small-cell neuroendocrine prostate cancer (tNEPC) associated genes (fold change ≥ 1.5, 330 genes) from the *Aggarwal et al* dataset (c) in naïve, shScr LTenza, and DKD LTenza 16D cells. Data represent mean +/- SEM. (d) GSEA of Beltran_NEPC_UP genes in DKD LTenza and shScr LTenza 16D cells. (e and f) GSEA of Hallmark_Myc_targets in DKD LTenza and naïve 16D cells (e), or DKD LTenza and shScr LTenza 16D cells (f). (g) Heatmap of select glycolytic genes from 3 technical replicates per line. (h) Western blot detecting DRP1 phosphorylation at S616 (DRP1-P616), DRP1, PSA, and Actin (control) in naïve, shScr LTenza, and DKD LTenza 16D lysates. (i-k) Maximal respiration (i), mitochondrial (Mito) ATP production (j), and glycolytic ATP production (k) in naïve, shScr LTenza, and DKD LTenza 16D cells from 4 biological replicate experiments. (l) Total ATP production, represented as the sum of mitochondrial ATP production (Mito ATP) and glycolytic ATP production (Glyco ATP), of naïve, shScr LTenza, and DKD LTenza 16D cells from 4 biological replicate experiments. Statistics refer to comparison of total ATP levels. Data represent mean +/- SEM. (m) Percentage of total ATP production from mitochondrial ATP production (% ATP from Mito) of naïve, shScr LTenza, and DKD LTenza 16D cells from 4 biological replicate experiments. (n) Relative sensitivity of the proliferation of naïve, shScr LTenza, and DKD LTenza 16D cells to 72-hour treatment with 30nM IACS. Significance was evaluated (d-f) using normalized enrichment scores (NES) and false discovery rates (FDR). P-values were calculated from an unpaired t-test with Welch's correction (a-c, l and n). $^{*}p < 0.05$, $^{**}p < 0.01$, $^{***}p < 0.001$, $^{****}p < 0.0001$, n.s. = not significant, $p \geq 0.05$.

Supplementary Figure 7

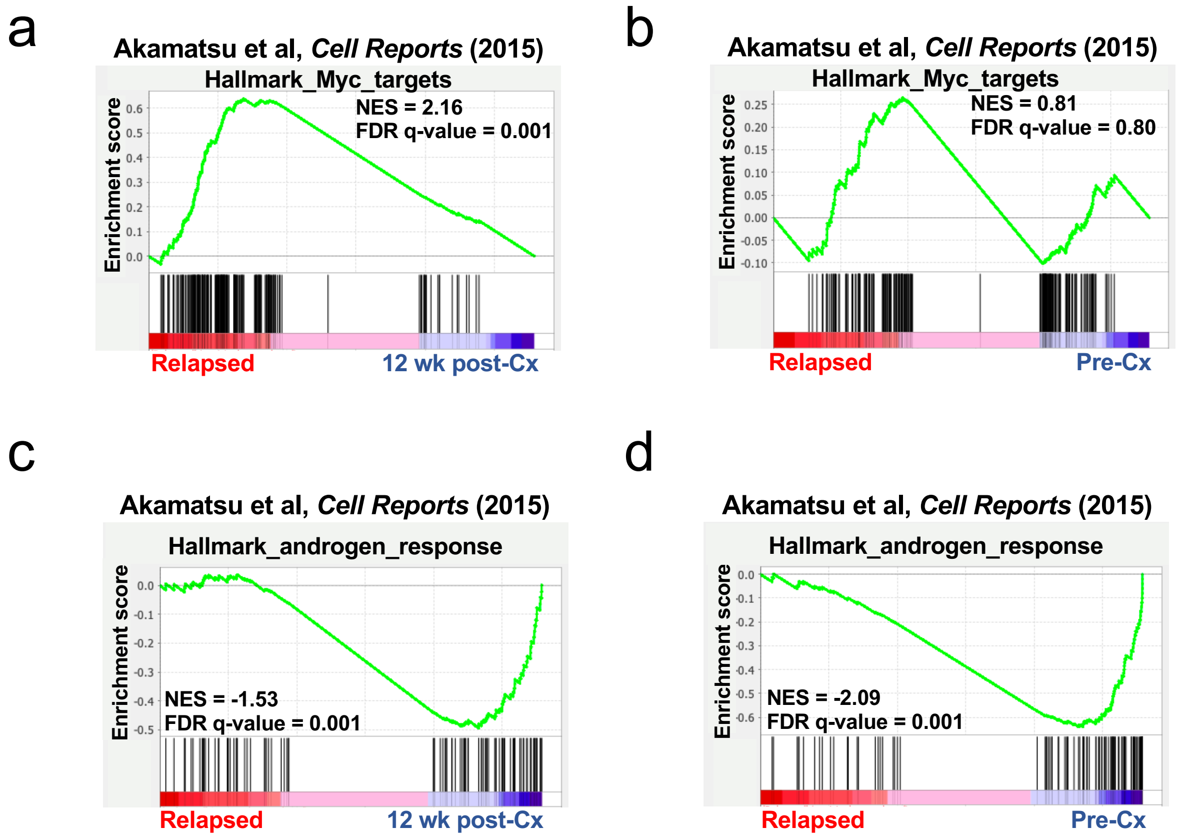

**Supplementary Figure 7. MYC target gene expression is restored in LTL331 model of relapsed castration-resistant prostate cancer despite maintenance of low AR activity.** (a and b) GSEA of Hallmark_Myc_targets in relapsed and 12-week post-castration (12 wk post-Cx) LTL331 tumor samples (a) and relapsed and pre-castration (Pre-Cx) samples (b) from the *Akamatsu et al* dataset. (c and d) GSEA of Hallmark_androgen_response genes in relapsed and 12 wk post-Cx LTL331 tumor samples (c) and relapsed and pre-Cx samples (d) from the *Akamatsu et al* dataset. Significance was evaluated (a-d) using normalized enrichment scores (NES) and false discovery rates (FDR).
